# Zona incerta dopamine neurons encode motivational vigor in food seeking

**DOI:** 10.1101/2023.06.29.547060

**Authors:** Qiying Ye, Jeremiah Nunez, Xiaobing Zhang

## Abstract

Energy deprivation triggers food seeking to ensure homeostatic consumption, but the neural coding of motivational vigor in food seeking during physical hunger remains unknown. Here, we report that ablation of dopamine (DA) neurons in zona incerta (ZI) but not ventral tegmental area potently impaired food seeking after fasting. ZI DA neurons were quickly activated for food approach but inhibited during food consumption. Chemogenetic manipulation of ZI DA neurons bidirectionally regulated feeding motivation to control meal frequency but not meal size for food intake. In addition, activation of ZI DA neurons and their projections to paraventricular thalamus transited positive-valence signals to promote acquisition and expression of contextual food memory. Together, these findings reveal that ZI DA neurons encode motivational vigor in food seeking for homeostatic eating.

**One Sentence Summary:** Activation of ZI DA neurons vigorously drives and maintains food-seeking behaviors to ensure food consumption triggered by energy deprivation through inhibitory DA^ZI-PVT^ transmissions that transit positive-valence signals associated with contextual food memory.

---

Daily food intake is well controlled by the brain to sense the body’s energy states and then drive food-seeking behaviors that ensure homeostatic food consumption. It is relatively clear that both the hypothalamic arcuate nucleus (ARC) and the nucleus of the solitary tract (NTS) sense the body’s energy states to initiate or terminate homeostatic eating. However, it remains unknown how both homeostatic and satiety signals are integrated by the brain to drive and maintain food-seeking behaviors particularly during energy deprivation. Mice with global dopamine (DA) deficiency were reported to be aphagic with severe starvation (*1*). However, activation of ventral tegmental area (VTA) DA neurons or their projections to nucleus accumbens only increased meal frequency but not cumulative food consumption (*2, 3*). Although VTA DA neurons may participate in some feeding-related behaviors especially hedonic eating, VTA DA signaling is not necessary for homeostatic eating especially food-seeking behaviors triggered by energy deficit (*4*). Therefore, it is critically important to decipher the neural coding of motivational vigor in food-seeking behaviors following central sensing of energy deficiency.

## DA neurons in the ZI but not the VTA are required for maintaining motivational vigor in fasting-triggered food seeking

To revisit a role of VTA DA neurons in regulating food motivation, we injected AAV into VTA of TH-Cre mice to express caspase (Casp) for ablating VTA DA neurons (fig. S1A and B). VTA DA ablation decreased daily meal numbers but increased meal size tested with feeding experimentation devices (FED) in home cages under free-feeding mode (fig. S1C) (*5*), but had little effect on daily food consumption and body weight gain (fig. S1D). Using operant behavioral tests under both fixed-ratio (FR) and progressive-ratio schedules of reinforcement (*6*), we found mice with VTA DA ablation had significantly lower active lever presses during both FR and PR sessions as well as breakpoints for food reward during PR sessions in fed but not fasted conditions (fig. S1E-H). This is consistent with previous findings that VTA DA neurons are involved in regulating motivation for palatable food intake in fed but not fasted states (*7-9*). To further reveal a source of DA neurons that are responsible for the DA control of homeostatic eating since mice with global DA knockout were aphagic (*1*). Our attention was drawn to A13 DA neurons in the ZI, an area we recently reported to regulate feeding and body weight gain (*10*). We similarly injected AAV to selectively ablating ZI DA neurons in TH-Cre mice (Fig. 1, A and B). ZI DA ablation reduced meal frequency without an effect on meal size to decrease daily food intake and body weight gain (Fig. 1, C and D). Furthermore, we found mice with ablation of ZI DA neurons had no effect on active lever presses and breakpoints for food during both FR and PR sessions in fed mice (Fig. 1E, and fig. S2B-D). However, compared to control, mice with ZI DA ablation had a significantly lower active lever presses for breakpoints to obtain food during PR sessions after fasting of 24 h (Fig. 1F and fig. S2E). ZI DA ablation also decreased active lever presses during reinstated PR sessions following extinction (Fig. 1G). In addition, mice with ZI DA ablation showed a decreased motivation for food in an anxiety-conflict condition with increased latency and decreased entries to the center of open-field chambers when food was placed in the center (Fig. 1, H and I, fig. S3). Together, these findings indicated that DA neurons in ZI but not VTA are required for driving vigorous effort in food-seeking behaviors that ensure subsequent food consumption during energy deficiency.

**Fig. 1.**
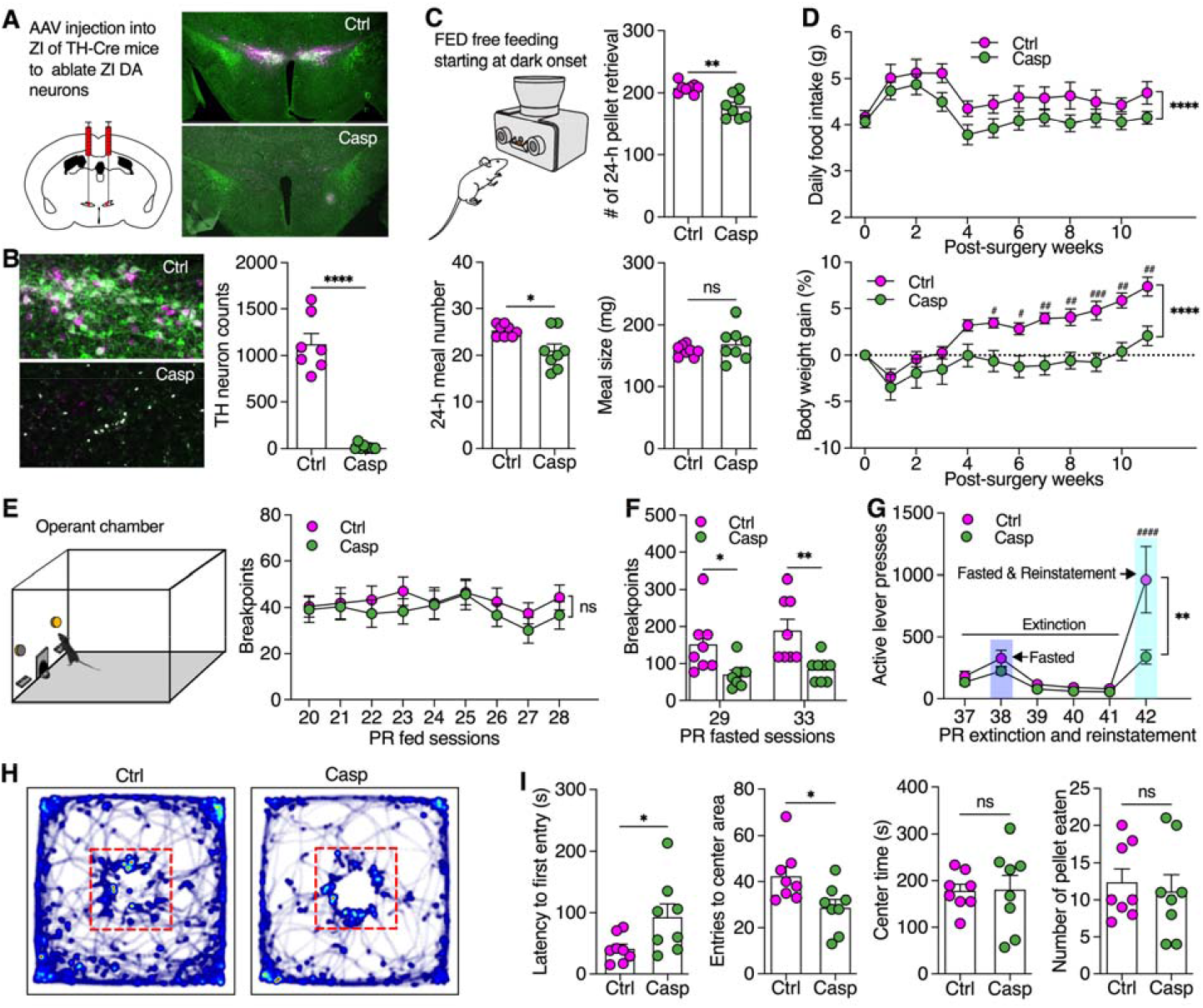
Selective ablation of ZI DA neurons impaired motivational vigor in fasting-triggered food seeking. (**A**) AAV was injected into bilateral ZI of TH-Cre mice to induce mCherry (n= 8 mice) or mCherry plus caspase expression (n=8 mice) in ZI DA neurons. (**B**) TH-immunoreactive ZI neurons were ablated by virus-induced caspase expression. (**C**) Meal pattern analysis using FED free feeding shows total number of pellet retrieval, meal numbers, and averaged meal size of 24 h. (**D**) Daily food intake and body weight gain in both control and caspase mice for 11 weeks following virus injection. (**E**) Breakpoint reached by fed mice during operant PR sessions of 45 min. (**F**) Breakpoints reached by 24 h fasted mice. (**G**) Active lever presses during PR extinction and reinstatement sessions. (**H**) Real-time activity of mice in open-field chambers with food pellets placed in the center. (**I**) The latency to the first entry of the center area, the total entries to center area, the total time in the center area, and the number of food pellets consumed by mice in open-field chambers with food pellets placed in the center. Unaired t test for (*B, C,I*), Two-way ANOVA with Post hoc Bonferroni test for (*D-G*).\**P* < 0.05, \*\**P* < 0.01, \*\*\**P* < 0.001, \*\*\*\**P* < 0.0001, ^####^*P* < 0.0001; ns: no significance.

### ZI DA neurons bidirectionally regulate motivated food seeking to tune meal frequency for daily feeding control

To determine how ZI DA neurons regulate feeding, we then injected AAV to express hM3D(Gq) selectively in ZI DA neurons of TH-Cre mice for chemogenetic activation (Fig. 2A, and fig. S4). The function of hM3D(Gq) was confirmed with CNO (3.0 μM) using slice patch-clamp recordings (Fig. 2B). Chemogenetic activation of ZI DA neurons with IP injection of CNO (2.0 mg/kg) in both sexes strongly increased regular food intake in both light and dark cycles as well as refeeding after 24 h fasting (Fig. 2C, and fig. S5, A and B). Electrophysiological data indicated that high-fat high-sucrose (HFHS) diet for two weeks depolarized the resting membrane potentials and increased the frequency of excitatory postsynaptic currents onto ZI DA neurons (fig. S6). Chemogenetic activation of these neurons also increased HFHS consumption in mice (fig. S5C). Furthermore, chemogenetic activation of ZI DA neurons potently promoted active lever presses to increase breakpoints for food reward of both light and dark cycles in both fed and fasted mice (Fig. 2D and fig. S7). However, CNO injection in control mice with mCherry expression in ZI DA neurons had no effect on food intake or operant lever presses during PR sessions (fig. S8). Using FED devices under both free-feeding mode and FR1 mode with nose pokes for food serving, we found chemogenetic activation of ZI DA neurons increased daily meal numbers, decreased inter-meal intervals, but did not affect meal size or meal duration (Fig. 2E and F, fig. S9). These results thus indicate that activation of ZI DA neurons promotes motivated effort to increase meal frequency for food intake especially in dark cycles and food-deprived conditions.

**Fig. 2.**
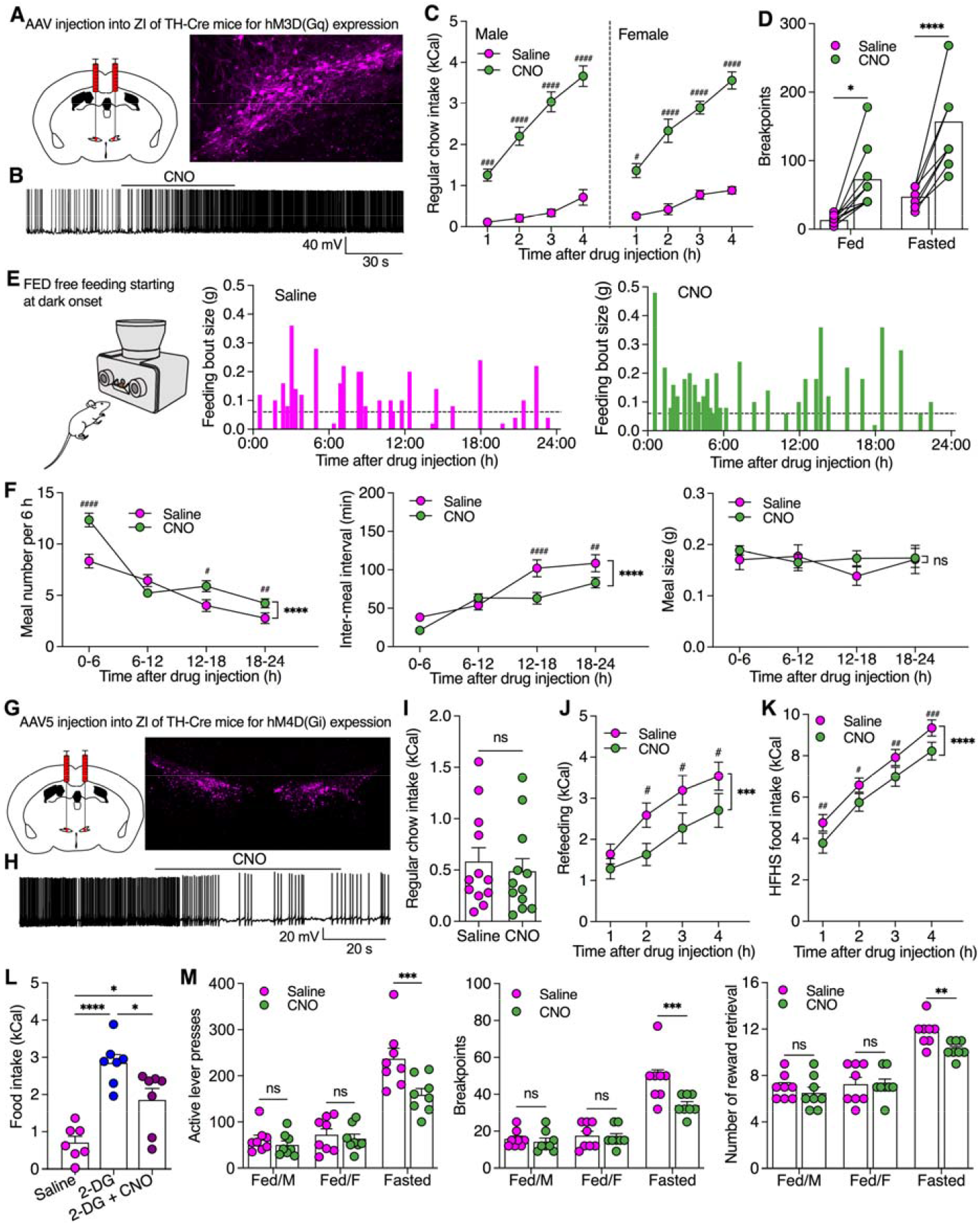
Chemogenetic activation of ZI DA neurons promoted, but inhibition of these neurons reduced, vigor in food seeking to regulate food consumption. (**A**) AAV was injected to bilateral ZI of TH-Cre mice to express excitatory hM3D(Gq) in ZI DA neurons. (**B**) A representative trace from slice recording shows that CNO (3 μM) excited a hM3D(Gq)-positive ZI DA neuron. (**C**) Regular food consumption during light cycle in both male (n= 12) and female (n= 8) TH-Cre mice with hM3D(Gq) expression in ZI DA neurons following IP injection of saline or CNO (2.0 mg/kg). (**D**) Breakpoints reached by both fed (n= 9) and partially fasted mice (70% of their daily food intake, n= 9) during PR sessions following IP injections of saline or CNO (2.0 mg/kg). (**E**) Real-time feeding bout sizes of a female ZI^TH-hM3D(Gq)^ mouse following IP injection of saline and CNO (2.0 mg/kg). FED devices at free-feeding modes were used for collecting real-time food intake in home cages. (**F**) Bar graphs showing meal numbers, inter-meal intervals, and meal sizes of 6 h. n= 9 female mice each group. (**G**) AAV was injected to bilateral ZI of TH-Cre mice to express inhibitory hM4D(Gi) in ZI DA neurons. (**H**) A representative trace shows that CNO (3.0 μM) inhibited a ZI^TH-hM4D(Gi)^ neuron. (**I**) Regular food intake of fed mice (n= 12) over 4 h following saline and CNO injection. (**J**) Refeeding after 24 h fasting following saline and CNO injection. (**K**) HFHS food intake following saline and CNO injection. (**L**) Food intake over 4 h following injection of saline, 2-DG (200 mg/kg), or 2-DG plus CNO (2.0 mg/kg). n = 7 each group. (**M**) Bar graphs showing active lever presses, breakpoints, and number of rewards earned in both fed and fasted mice (70% of their daily food intake overnight) during operant PR tests of 45 min following saline or CNO (2.0 mg/kg) injection. Three-way ANOVA with Post hoc Bonferroni test for (*C*), two-way RM ANOVA with Post hoc Bonferroni test (*D, F, J, K, M*), paired t test for (*I*), one-way ANOVA with Post hoc Bonferroni test (*L*), \**P* < 0.05, \*\**P* < 0.01, \*\*\**P* < 0.001, \*\*\*\**P* < 0.0001; ^#^*P* < 0.05, ^##^*P* < 0.01, ^###^*P* < 0.001, ^####^*P* < 0.0001; ns: no significance.

We further injected AAV to express hM4D(Gi) in ZI DA neurons of TH-Cre mice for chemogenetic inhibition (Fig. 2G). CNO (3.0 μM) hyperpolarized hM4D(Gi)-positive ZI DA neurons to decrease the neuronal activity (Fig. 2H). Chemogenetic inhibition of ZI DA neurons had little effect on regular food intake in fed conditions but decreased refeeding after 24 h fasting as well as HFHS food intake (Fig. 2I-K). In addition, chemogenetic inhibition of ZI DA neurons impaired glucoprivic feeding evoked by IP injection of 2-Deoxy-D-glucose (2-DG) (*11, 12*) (Fig. 2L). Furthermore, we found chemogenetic inhibition of ZI DA neurons decreased active lever presses to reduce breakpoints for food reward primarily in fasted but not fed mice (Fig. 2M). As a result, chemogenetic inhibition of ZI DA neurons decreased daily meal numbers but not meal size in mice (fig. S10). Together, these data indicate that inhibition of ZI DA neurons decreases motivational effort in food seeking to reduce food consumption selectively in fasted conditions, suggesting ZI DA neurons participate in physiological feeding triggered by energy deficit.

### ZI DA neurons promote both acquisition and expression of contextual memory for food seeking

To determine whether ZI DA neurons drive food-seeking behaviors through promoting contextual reward memory, we tested operant behaviors during post-training sessions without food delivery (Fig. 3A). Following the operant trainings, mice learned to press levers to obtain food reward. Although no pellet was delivered after lever presses during post-training sessions, mice still compulsively pressed the lever that was paired to food reward previously. Chemogenetic activation of ZI DA neurons strongly increased compulsive lever presses for food seeking (Fig. 3B-D).

**Fig. 3.**
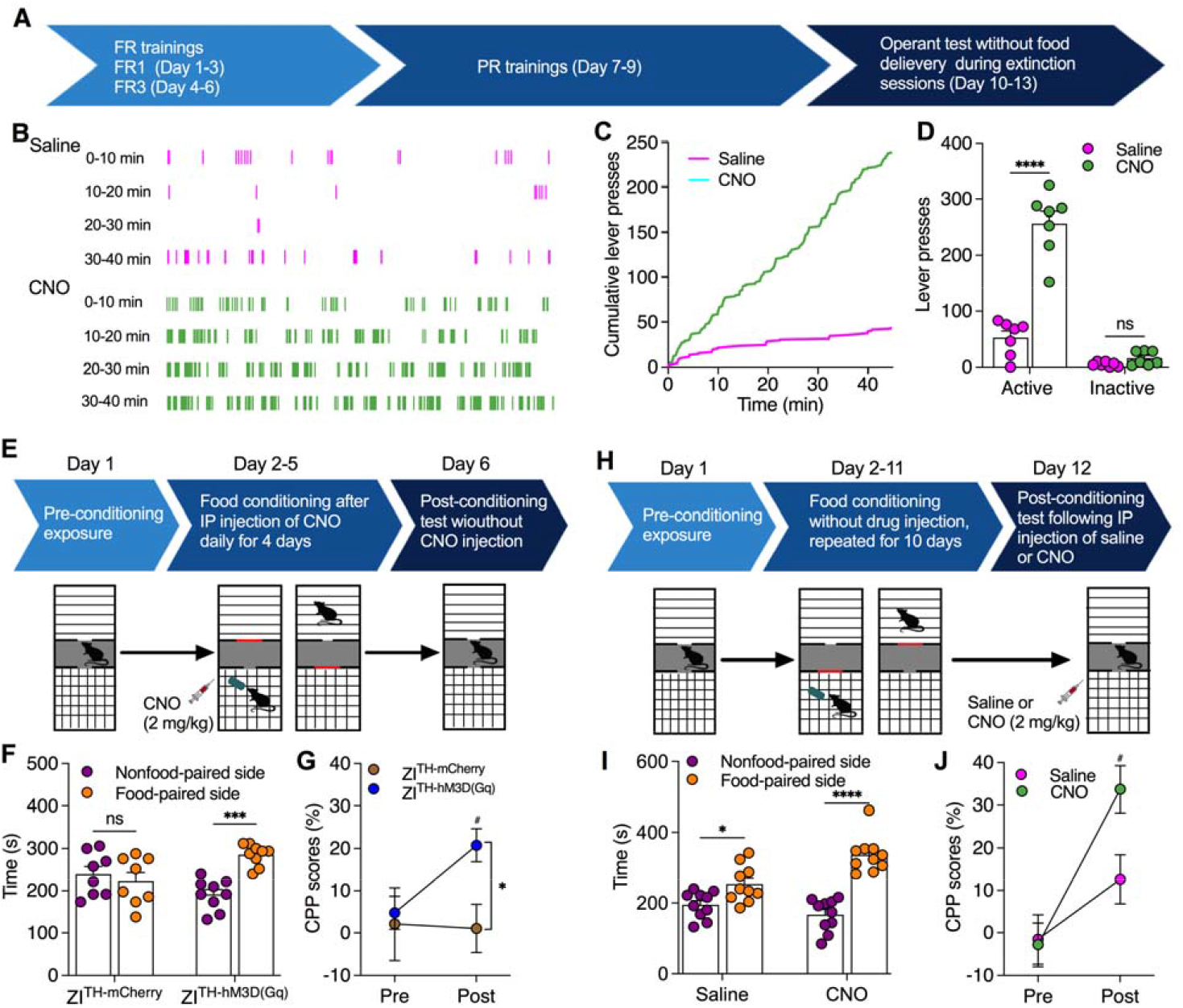
Activation of ZI DA neurons promoted both acquisition and expression of contextual food memory. (**A**) Timeline for operant training and tests. (**B**) Real-time lever presses during operant tests without reward delivery. (**C**) Cumulative lever presses during operant tests without reward delivery. (**D**) Presses of the food-paired active levers and the nonfood-paired inactive levers during operant tests without reward delivery. n= 7 mice each group. **(E**) Timeline for testing food CPP acquisition in both ZI^TH-mCherry^ (n= 8) or ZI^TH-hM3D(Gq)^ (n= 9) mice. (**F**) Time that mice spent in both nonfood-paired and food-paired sides during post-conditioning sessions. (**G**) CPP scores during both pre-conditioning and post-conditioning tests. (**H**) Timeline for testing food CPP expression. ZI^TH-hM3D(Gq)^ mice received 10-day place conditioning with HFHS food as well as pre- and post-conditioning place preference tests. (**I**) Time that mice spent in both nonfood-paired and food-paired sides during post-conditioning sessions following saline (n= 10 mice) or CNO (2.0 mg/kg, n= 10 mice) injection. (**J**) CPP scores in mice during both pre-conditioning and post-conditioning tests. Two-way RM ANOVA with Post hoc Bonferroni test for (*D*, G, *J*). Two-way ANOVA with Post hoc Bonferroni test for (*F, I*). \**P* < 0.05, \*\*\**P* < 0.001, \*\*\*\**P* < 0.0001; ^#^*P* < 0.05; ns: no significance.

We then asked whether ZI DA neurons regulate contextual memory associated with food reward for food seeking, we performed a conditioned place preference (CPP) paradigm to test the preference to food-conditioned environment. Control ZI^TH-mCherry^ mice didn’t develop a preference for food-conditioned chambers after 4-day trainings when intraperitoneal CNO (2 mg/kg) injection was given 30 min before food conditioning daily (Fig. 3E-G). However, ZI^TH-hM3D(Gq)^ mice with CNO injection preferred food-paired chambers after 4-day food-paired trainings (Fig. 3E-G). These data suggest that chemogenetic activation of ZI DA neurons promote acquisition of contextual food memory. We further used ZI^TH-hM3D(Gq)^ mice with longer food-paired trainings for 10 days to test a role of ZI DA neurons in expression of contextual food memory. All mice developed preferences for food-conditioned chambers after 10-day food-paired trainings (Fig. 3H-J), while chemogenetic activation of ZI DA neurons during post-conditioning sessions significantly increased the preference for food-conditioned chambers (Fig. 3H-J). These data thus suggest activation of ZI DA neurons also promote expression of contextual memory for food seeking, which is supported by previous findings that ZI integrates multiple sensory information for potential motivational drive(*13*).

### A distinct role of ZI DA neurons in feeding control from ZI GABA neurons

Early studies have reported that sheep ZI neurons were activated responding to the approach of food (*14, 15*). Neurons in the rostromedial ZI, the location of A13 DA neurons, were also activated by food teasing but not refeeding in fasted mice (*16*). These studies together suggest that ZI neurons are activated by food-associated signals to motivate food-seeking for food intake. To determine whether and how ZI DA neurons participate in the physiological feeding control, we applied in vivo fiber photometry to dynamically monitor the real-time activity of ZI DA neurons with AAV-induced GCaMP7f expression under a free-feeding test using FED devices (Fig. 4A). We found ZI DA neurons quickly increased their activity for approaching food and immediately decreased the activity after pellet retrieval followed by a rapid inhibition during food consumption (Fig. 4B and C) before it went back to baseline level. Using a similar approach, we also recorded a response of ZI GABA neurons (Fig. 4A), the neighboring neurons that we previously reported to induce binge-like eating(*10*), to feeding-related behaviors. In contrast to ZI DA neurons, ZI GABA neurons had a relatively smaller increase in activity while approaching food but a larger and long-lasting suppression during food consumption (Fig. 4D-F). The activity response of ZI DA neurons is also different from that of ARC AgRP neurons which was constantly inhibited by food consumption (*17*). Together, these data indicate that both ZI DA and GABA neurons physiologically participate in feeding regulation. However, ZI DA neurons are more active in response to food approach, suggesting ZI DA neurons regulate food intake differentially from ZI GABA neurons as well as ARC AgRP neurons.

Using both RNAScope in situ hybridization, transgenic GAD67-GFP mice, and TH immunocytochemistry, we found ZI TH neurons co-expressed GAD2/GAD65 but not GAD1/GAD67 (fig. S11A, S11B, and S12A). This is consistent with a previous study using TH and GAD double immunostaining that showed little co-expression in the ZI (*18*). To further understand a difference of ZI DA and GABA neurons in regulating feeding behavior for food intake, we injected AAV to express hM3D(Gq) in ZI DA neurons of TH-Cre mice and GABA neurons of vGAT-Cre mice for operant PR tests to compare the effects of the two types of ZI neurons on motivated food seeking (Fig. 4G and H). Only a small population of TH neurons were detected with vGAT-mCherry in rostrolateral ZI of vGAT-Cre mice with AAV injection (3.7%) and sparse vGAT mRNA expression (Fig. 4H and I, fig. S11C, fig. S13). Surprisingly, chemogenetic activation of ZI GABA neurons promoted active lever presses to increase breakpoints (BP) for food reward in only 8 out of 12 vGAT-Cre mice but decreased active lever presses in other 4 mice during PR sessions (Fig. 4J). However, chemogenetic activation of ZI GABA neurons increased food intake in both BP-increased and BP-decreased mice when food was freely available (Fig. 4K), suggesting differential neural coding for food seeking versus consumption. We further compared the effect of ZI GABA and DA neurons in the regulation of both food seeking and consumption. Chemogenetic activation of ZI GABA neurons had a comparably larger increase in food intake when food was freely available (Fig. 4L), while activation of ZI DA neurons had a significantly stronger increase in active lever presses and breakpoints for seeking food reward tested during PR sessions (Fig. 4M).

**Fig. 4.**
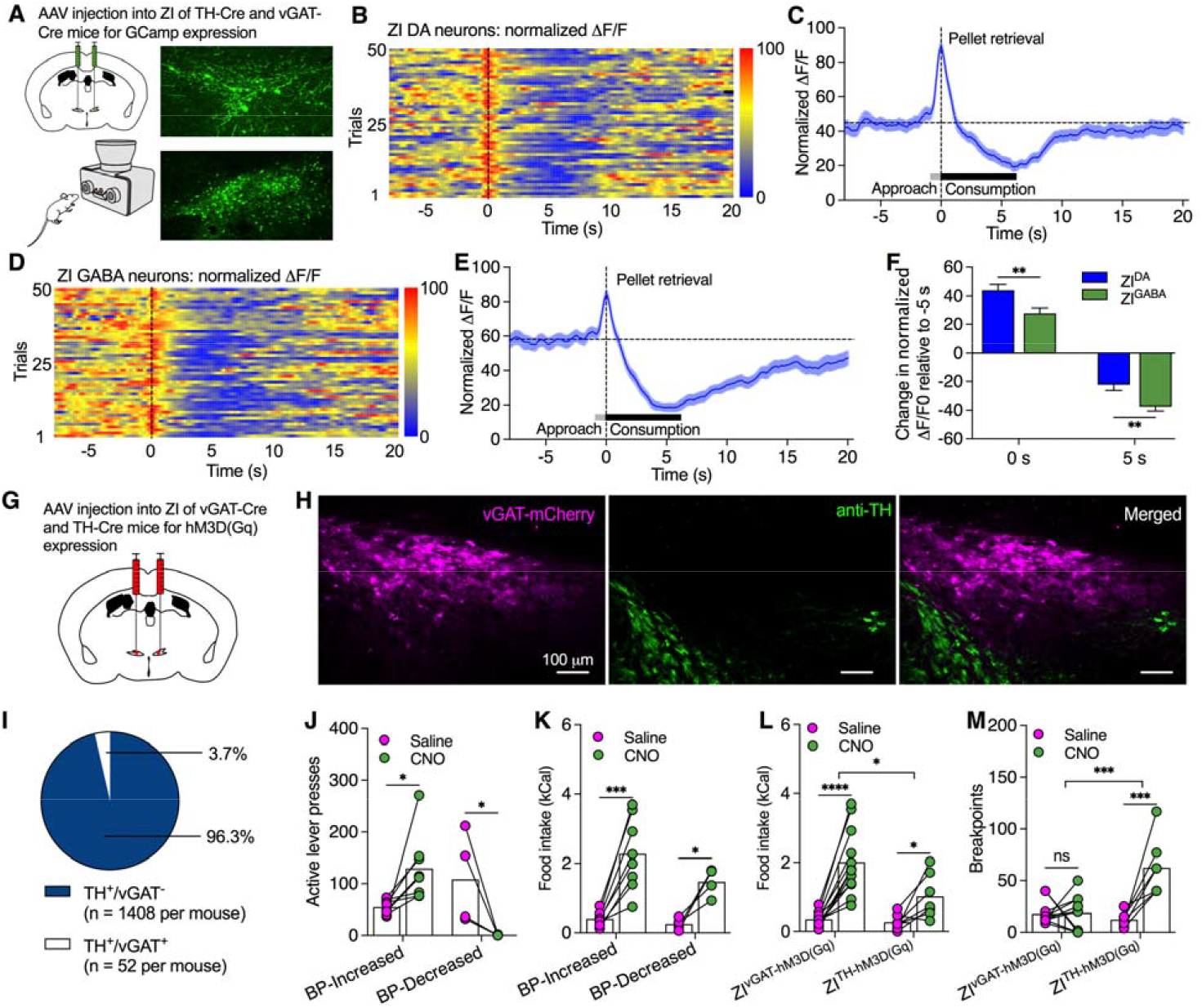
ZI DA and GABA neurons differentially regulate both food seeking and consumption. **(A**) AAV was injected into ZI of TH-Cre or vGAT-Cre mice to induce GcAMP expression in ZI DA or GABA neurons. (**B**) Normalized ΔF/F of ZI DA neurons from 50 feeding trials using FED devices at free-feeding modes. (**C**) Normalized ΔF/F of ZI DA neurons aligned to pellet retrieval from 50 feeding trials. (**D**) Normalized ΔF/F of ZI GABA neurons from 50 feeding trials. (**E**) Normalized ΔF/F of ZI GABA neurons aligned to pellet retrieval from 50 feeding trials. (**F**) Change in ΔF/F of ZI DA and GABA neurons for food approaching (0 s) and consumption (5 s). n= 50 trials from 4 mice for each group. (**G**) AAV was injected into bilateral ZI of vGAT-Cre and TH-Cre mice. (**H**) ZI^vGAT-hM3D(Gq)^ neurons were not colocalized with TH. (**I**) Percentages of both TH^+^/vGAT^-^ and TH^+^/vGAT^+^ neurons in the ZI. (**J**) Chemogenetic activation of ZI^vGAT-hM3D(Gq)^ neurons increased active lever presses in a breakpoint (BP)-increased group (n= 8 mice) but decreased in another BP-decreased group (n= 4 mice) during PR sessions. (**K**) Chemogenetic activation of ZI^vGAT-hM3D(Gq)^ neurons increased food intake in both BP-Increased (n= 8) and BP-Decreased mice (n= 4). (**L**) Comparison of food intake of both ZI^vGAT-hM3D(Gq)^ (n= 12) and ZI^TH-hM3D(Gq)^ (n= 7) mice. **(M**) Comparison of breakpoints reached by ZI^vGAT-hM3D(Gq)^ and ZI^TH-hM3D(Gq)^ mice. Two-way RM ANOVA with Post hoc Bonferroni test for (*F,J-M*). \**P* < 0.05, \*\*\**P* < 0.001, \*\*\*\**P* < 0.0001. ns: no significance.

As previously reported with optogenetic activation (*10*), we observed licking and gnawing-like behaviors following chemogenetic activation of ZI GABA neurons. This is also similar to activation of lateral hypothalamic (LH) GABA neurons and their projections to VTA that had no effect on food seeking but produced gnawing behavior (*19-21*). However, we didn’t observe any gnawing behavior by chemogenetic activation of ZI DA neurons, suggesting ZI DA neurons have little direct control of consummatory behavior but indirectly regulate food intake through promoting motivated food seeking.

### ZI DA neurons transit a positive-valence signal through inhibitory DA transmissions to PVT for feeding regulation

ZI GABA neurons send dense projections to PVT, an area that regulates behavioral motivation, for feeding control (*10, 22-25*). Inactivation of PVT neurons also promoted contextual place preference for sucrose seeking (*26*). To examine whether ZI DA neurons project to PVT for feeding regulation, we injected AAV to express EGFP in TH-positive ZI DA neurons of TH-Cre mice (Fig. 5A). EGFP-positive axons were detected in PVT, periaqueductal gray (PAG), and other areas (Fig. 4A, fig. S14). To further confirm ZI DA projections to PVT, we injected retrograde AAV into PVT of TH-Cre mice and found retrogradely labelled neurons in ZI (Fig. 5B). To identify a functional regulation of PVT neurons by ZI DA projections, we injected AAV to express ChIEF in bilateral ZI DA neurons of TH-Cre mice for slice recordings and behavioral tests with photostimulation of ChIEF-positive projections in PVT (Fig. 5C). Both photostimulation (20 Hz) and DA (30 μM) hyperpolarized and inhibited PVT neurons innervated by ChIEF-positive ZI DA terminals in the absence and presence of Bic (Fig. 5D, fig. S15A-C). The outward currents evoked by photostimulation of ZI-PVT DA projections were blocked by D2 receptor antagonist sulpiride (fig. S15D), suggesting postsynaptic D2 receptors in PVT neurons contribute to the inhibitory regulation of ZI-PVT DA pathways.

**Fig. 5.**
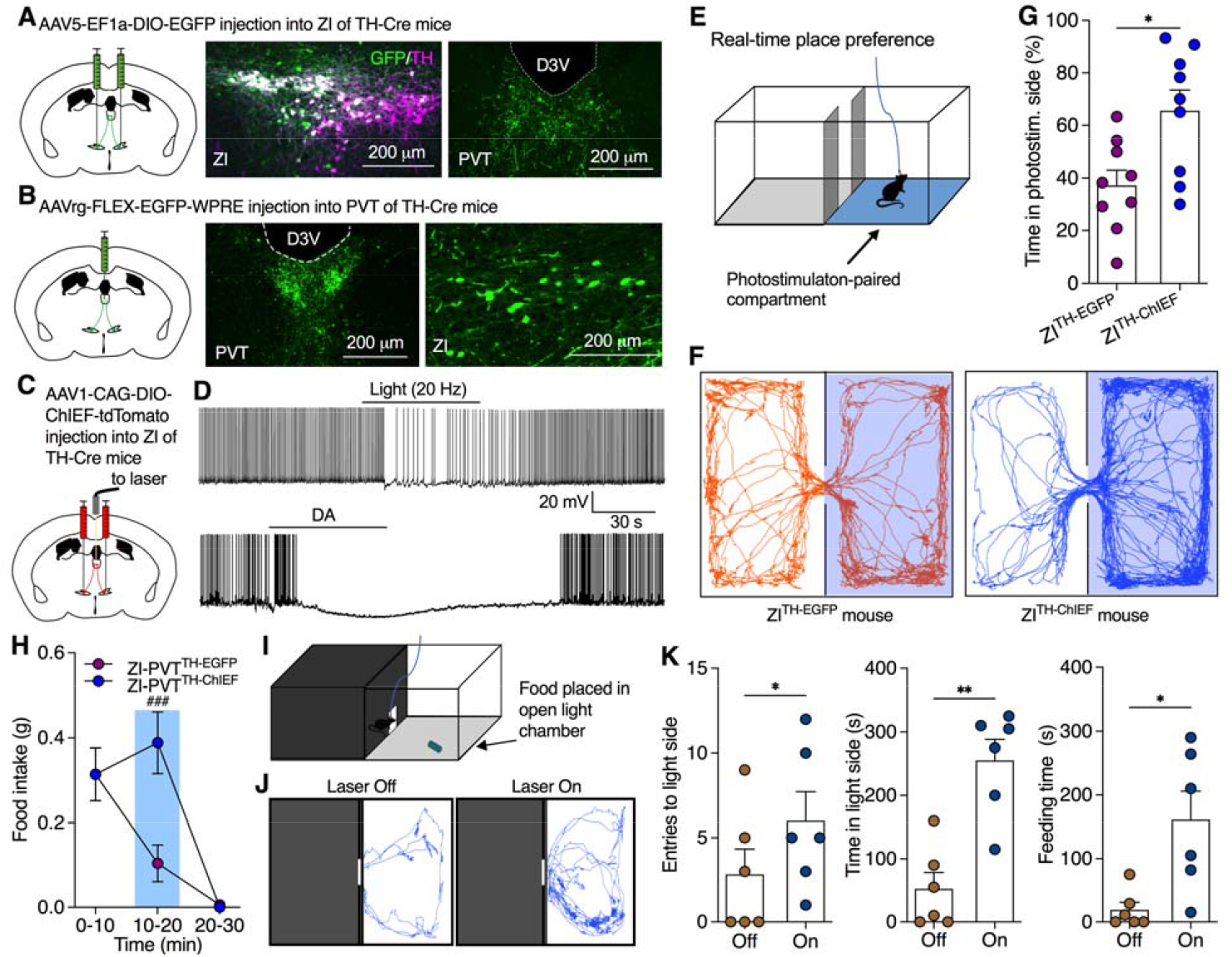
Optogenetic activation of ZI-PVT DA projections produced positive valence to promote food seeking for consumption. (**A**) AAV was injected into bilateral ZI of TH-Cre mice to induce EGFP expression in ZI DA neurons that sent EGFP-positive axons to PVT. (**B**) Retrograde AAV was injected into PVT of TH-Cre mice to trace ZI DA neurons that projected to PVT. (**C**) AAV was injected into bilateral ZI of TH-Cre mice to induce ChIEF or EGFP expression in ZI DA neurons. (**D**) Both photostimulation and DA (30 μM) inhibited PVT neurons surrounded by ChIEF-positive ZI DA axons. (**E**) A two-compartment chamber was used for real-time place preference test with one compartment paired with photostimulation (20 Hz). The laser was only turned on when mice entered the photostimulation-paired side. (**F**) Real-time motion of a control ZI^TH-EGFP^ and a ZI^TH-ChIEF^ mouse in the two-compartment chamber. (**G**) Percentage of time that mice spent in photostimulation-paired compartment. n= 9 mice each group. (**H**) HFHS food intake of mice before, during, and after PVT photostimulation. n= 7 mice each group. (**I**) A light/dark box with entrance connecting the two sides for food intake test over 10 min when food pellets were placed in the light side. (**J**) Real-time activity tracking in light side of the chambers during 10-min food intake test with or without photostimulation (20 Hz) of DA^ZI-PVT^ pathway. (**K**) Total entries to light side, cumulative time in light side, and cumulative feeding time during 10-min food intake. n= 6 mice per group. Unpaired t test for (*G*), two-way ANOVA with Post hoc Bonferroni test for (*H*), and paired t test for (*K*). \**P* < 0.05, \*\**P* < 0.01; ^###^*P* < 0.001.

Activation of both LH GABA neurons and VTA DA neurons was reported to produce positive valence for the regulation of food reward (*3, 27*). Using self-photostimulation of ZI-PVT DA projections in two-compartment chambers for real-time place preference tests (Fig. 5E), we found ZI^TH-ChIEF^ but not control ZI^TH-EGFP^ mice spent significantly longer time in photostimulation-paired side (Fig. 5F and G) and continuously press levers to obtain self-stimulation of ZI-PVT DA pathways in operant chambers (fig. S16A-D). ZI^TH-ChIEF^ mice still compulsively pressed the lever during the first two post-pairing sessions without photostimulation following daily photostimulation-paired sessions of 5 days (fig. S16D). To further determine if activation of ZI-PVT DA projections promotes positive valence for contextual rewarding memory, we tested photostimulation-evoked CPP expression (fig. S16E and F). Before photostimulation pairing, mice spent less time in the compartments with smooth floor. However, mice developed and maintained a preference for smooth-floor compartments that were paired with photostimulation during the training sessions (fig. S16G and H). After photostimulation-paired trainings for 3 days, mice still spent more time in photostimulation-paired compartments during post-pairing sessions (fig. S16G and H). Together, these results indicate that activation of ZI-PVT DA projections produces positive valence and promotes contextual memory for reward seeking.

Both optogenetic and chemogenetic activation of ZI DA axons locally in PVT increased food intake (Fig. 5H and fig. S17), while photostimulation of ZI-PAG DA projections produced little effect on both food intake and motivated food seeking (fig. S18). Light/dark boxes were previously used for testing feeding motivation in an anxiety-conflict environment (Fig. 5I) (*10*). Photostimulation of ZI-PVT DA projections significantly increased total entries to light compartment, total time spent in light side, and total feeding time (Fig. 5K). These results thus indicate that ZI-PVT DA activation promotes feeding motivation to increase food intake even in anxiety-associated conditions.

## Discussion

In conclusion, we have shown that ZI DA neurons encode motivational vigor in food-seeking behaviors triggered by energy deficiency. As a result, ZI DA neurons regulate meal frequency but not meal size to control food intake. PVT is a postsynaptic target for ZI DA neurons to transit positive valence which is activated to promote contextual memory for food seeking. This is different from AgRP neurons that transit a negative-valence signal for energy sensing which is modulated by food consumption and sensory food cues (*17, 28*). In contrast to aphagia and severe starvation caused by loss of AgRP neurons (*29*), ablation of ZI DA neurons selectively impaired motivational vigor in food seeking to decrease meal frequency that reduce daily food intake and weight gain (Fig. 1). Activation of ZI DA neurons promoted food-seeking behaviors in both fed and fasted mice, while activating AgRP neurons had no effect on food seeking in fasted mice (*30*). These differences thus suggest that AgRP neurons are constantly activated by fasting for energy sensing but ZI DA neurons are only activated by environmental food cues for food seeking.

The present study also reveals a critical role of ZI DA neurons distinctly from VTA DA neurons in regulating feeding motivation. Activation of VTA DA neurons or their projections to nucleus accumbens had little effect on cumulative food intake though it increased meal frequency and promoted food motivation in fed animals (*2, 3*). Consistent with these reports, our present data show that ablation of VTA DA neurons only reduced motivational vigor in food seeking in fed but not fasted mice. However, activation of ZI DA neurons increased feeding motivation in both fed and fasted mice, while ablation and silencing of these neurons reduced motivated food seeking selectively after fasting. The present findings thus demonstrate that ZI DA neurons drive food-seeking behaviors for homeostatic eating probably following metabolic sensing by other homeostasis-controlling neurons. In contrast, VTA DA neurons mainly regulate food reward for hedonic eating independent of physiological energy states. Together, these findings suggest that the brain uses two different DA pathways to drive hedonic and homeostatic eating.

ZI DA neurons are also distinct from LH GABA neurons which are activated for appetitive or consummatory behaviors (*19*). Activation of ZI DA neurons strongly promoted motivated food seeking and consumption but not gnawing behaviors. However, activation of LH GABA neurons induced gnawing behaviors for food consumption (*20, 21*). Activation of GABA^LH-VTA^ projections also induced compulsive eating but not reward seeking, while inhibition of these pathways did not reduce food intake in food-restricted mice (*21*). Similar to LH GABA neurons, ZI GABA neurons primarily regulate food consumption supported by the present data that activation of these neurons evoked gnawing behaviors for a striking increase in food consumption but slightly increased food seeking (*10*). Together, these findings suggest that ZI DA neurons are responsible for driving and maintaining food-seeking behaviors, which is different from both ZI and LH GABA neurons for their role in driving consummatory behavior.

## Acknowledgements

The authors thank Y. Yang and L. Zhang of Yale University for their technical supports in TH and DA immunocytochemistry, L. Barrett for some of the stereotactic surgery and behavioral tests, and D. Williams for providing us with their fluorescent microscope in the early stages of the project. We are grateful to Z. Wang for his comments on the manuscript. We also would like to thank the late A. van den Pol of Yale University for his support in initiating this project.

## Author contributions

X.Z. conceived and designed the experiments. Q.Y. and J.N. performed immunocytochemistry, stereotaxic surgeries for virus injections and fiber optic implantation, and behavior experiments. Q.Y and X.Z. performed slice electrophysiology and fiber photometry experiments. X.Z wrote the original draft. Q.Y and J.N. reviewed and edited the manuscript.

## Data availability

All materials and data are available from the corresponding author upon request.

## Supplementary Materials

Materials and Methods

References (31-35)

Fig. S1 to S18

