## Supplemental material for "Zona incerta dopamine neurons encode motivational vigor in food seeking"

### Supplementary Materials for

#### Zona incerta dopamine neurons encode motivational vigor in food seeking for homeostatic eating

Qiying Ye *et al*.

**The PDF file includes:**

Materials and Methods
References 31-35
Fig. S1- S18 and legends

##### Methods

##### Animals

Both TH-Cre and vGAT-Cre mice were obtained from the Jackson Laboratory (Bar Harbor, ME, USA). GAD67-GFP mice were provided by Pradeep G Bhide lab at Florida State University (*31*). TH-tdTomato mice were generated by Gensat/Rockefeller University and provided by Anthony van den Pol lab at Yale University (*32*). All mice were housed in a climate-controlled vivarium on a 12:12 h light/dark cycle and *ad libitum* access to food and water. All animals and experimental procedures in this study were approved by the Florida State University Institutional Animal Care and Use Committee. Both male and female mice used for this study were 8-10 weeks old at the beginning of the experiments.

##### Intracerebral viral vector injections and optic fiber implantation

For chemogenetic activation or inhibition of ZI DA neurons, AAV5-hSyn-DIO-hM3D(Gq)-mCherry (Addgene, MA, USA), AAV5-hSyn-DIO-hM4D(Gi)-mCherry (Addgene, MA, USA), or AAV5-hSyn-DIO-mCherry (Addgene, MA, USA) was injected into bilateral ZI (0.2 μl per side) of TH-Cre mice. The stereotaxic coordinates for viral injections from Bregma were: anterior-posterior (AP)= -1.3 mm, medial-lateral (ML)= +/-0.5 mm, dorsalventral (DV)= -4.5 mm from surface level of Bregma. One month after viral injection, mice were used to for behavioral tests with chemogenetic modulation.

For chemogenetic activation of ZI GABA neurons, AAV5-hSyn-DIO-hM3D(Gq)-mCherry (Addgene, MA, USA), or AAV5-hSyn-DIO-mCherry (Addgene, MA, USA) was injected into bilateral ZI (0.2 μl per side) of vGAT-Cre mice. The stereotaxic coordinates for viral injections from Bregma were: AP= -1.0 mm, ML= +/-0.7 mm, DV= -4.5 mm from surface level of Bregma. One month after viral injection, mice were used for behavioral tests with chemogenetic activation.

For optogenetic activation of ZI-PVT or ZI-PAG DA projections, AAV1-CAG-DIO-ChIEF-tdTomato (Signagen, MD, USA), AAV1-EF1a-DIO-ChR2(H134R)-EYFP-WPRE-HGHpA (Addgene, MA, USA), or AAV5-EF1a-DIO-EGFP (Addgene, MA, USA) was injected into bilateral ZI (0.2 μl per side) of TH-Cre mice. The stereotaxic coordinates for viral injections from Bregma were: AP= -1.3 mm, ML= 0.5 mm, DV= -4.5 mm from surface level of Bregma. The optic fiber (OD: 200 μm, NA: 0.22, Doric Lenses, Canada) was implanted to target PVT or PAG. The stereotaxic coordinates for PVT were: AP= -1.3 mm, ML= 0.05 mm, DV= -2.8 mm from surface level of Bregma. The stereotaxic coordinates for PAG were: AP= -4.8 mm, ML= +/-0.5 mm, DV= -2.0 mm from surface level of Bregma. One month after viral injection, mice were used for behavioral tests with photostimulation of PVT or PAG.

For fiber photometry recording of ZI DA and GABA neurons, AAV5-syn-FLEX-jGCaMP7f-WPRE (Addgene, MA, USA) was injected into ZI (0.2 μl) of TH-Cre mice and vGAT-Cre mice. The stereotaxic coordinates for viral injections from Bregma of TH-Cre mice were: AP= -1.3 mm, ML= 0.5 mm, DV= -4.5 mm from surface level of Bregma. The optic fiber was implanted to target ZI of injection side. The stereotaxic coordinates for optic fiber implant from Bregma were: AP= -1.3 mm, ML= 0.5 mm, DV= -4.5 mm from surface level of Bregma. The stereotaxic coordinates for viral injections from Bregma of vGAT-Cre mice were: AP= -1.3 mm, ML= 0.7 mm, DV= -4.5 mm from surface level of Bregma. The optic fiber was also implanted to target ZI of injection side. The stereotaxic coordinates for optic fiber implant from Bregma were: AP= -1.0 mm, ML= 0.7 mm, DV= -4.3 mm from surface level of Bregma. One month after viral injection, mice were used for photometry tests.

For ablation of ZI and VTA DA neurons, 200 nl AAV5-hSyn-DIO-mCherry (1:5 dilution in PBS, control group) or AAV5-flex-taCasp3-TEVp mixed with AAV5-hSyn-DIO-mCherry (1:5 dilution in AAV5-flex-taCasp3-TEVp, Casp ablation group) was injected into bilateral ZI (coordinates: AP= -1.3 mm, ML= +/-0.5 mm, DV= -4.5 mm) or VTA (coordinates: AP= -3.1 mm, ML= +/-0.4 mm, DV= -4.5 mm) of TH-Cre mice. Mice were measured about daily food intake and weekly body weight before, during, and after injection surgery.

**Slice preparation and patch-clamp electrophysiology**

Both male and female mice were used for preparing coronal brain slices (300 μm thick) containing the PVT as detailed in our previous studies(*10, 32*). For whole-cell recording of PVT neurons, fresh brain slices were transferred to a recording chamber mounted on a Zeiss upright microscope (Zeiss, Berlin, Germany) and perfused with a continuous flow of gassed ACSF solution containing (in mM) 124 NaCl, 3 KCl, 2 $MgCl_2$, 2 $CaCl_2$, 1.23 $NaH_2PO_4$, 26 $NaHCO_3$, and 10 glucose (gassed with 95% $O_2$/5% $CO_2$; 300–305 mOsm). Pipettes used for whole-cell recording had resistances ranging from 4 to 7 MΩ when filled with pipette solution containing (in mM) 145 potassium gluconate, 1 $MgCl_2$, 10 HEPES, 1.1 EGTA, 2 Mg-ATP, 0.5 $Na_2$-GTP, and 5 disodium phosphocreatine (pH 7.3 with KOH; 290–295 mOsm). The recording was performed at 33 ± 1°C using a dual-channel heat controller (Warner Instruments, Holliston, MA, USA). EPC-10 USB amplifier (HEKA Instruments, NY, USA) and PatchMaster 2x90.5 software (HEKA Elektronik, Lambrecht/Pfalz, Germany) were used to acquire and analyze the data. For voltage-clamp recording, the membrane potentials were held at -70 mV. Traces were processed using Igor Pro 6.37 (Wavemetrics, OR, USA). Spontaneous postsynaptic currents were analyzed with MiniAnalysis 6.03 (Synaptosoft Inc., GA, USA).

**Operant conditioning and progressive ratio schedules of reinforcement**

Before operant conditioning training in mouse operant chambers (Med Associates, VT, USA), all C57BL/6J mice were food-restricted (70% of their daily food intake) to facilitate the acquisition of lever-press responding until they learned to press the lever to obtain the food pellet in 3 to 5 days. Mice were provided their daily quota of food in the home cage after the termination of the training session. We used a program designed by our programing engineer for the data acquisition. During the training, mice were initially trained under fixed-ratio 1 (FR1) sessions for 45 min daily. Animals had a choice between two press levers, an active lever press associated with a 3 sec light cue, and a concomitant delivery of high-fat high-sucrose (HFHS) pellets (20 mg, 48.9% Kcal as fat, Bio-Serv, NJ, USA) and an inactive lever press that remained inoperative and served as a control for general activity. Each active lever press triggered the delivery of one pellet during FR1 sessions. The lever remained inactive for 5 sec after each food delivery so that mice were able to retrieve the pellet and supplementary presses during the inactive period did not drive food delivery. After a training period of about 7-10 days when three successive sessions of obtaining equal and more than 20 pellets during the FR1 session of 45 min, mice were then engaged in consecutive PR sessions of 45 min. For the PR session, the number of lever presses required for one food pellet delivery followed the order (calculated by the formula [$5e^{(R*0.2)}$] - 5 where $R$ is equal to the number of food rewards already earned plus 1): 1, 2, 4, 6, 9, 12, 15, 20, and so on(*6*). The maximal number of active lever presses performed to reach the final ratio was defined as the breakpoint, a value reflecting animals' motivation to get the food reward.

**Food intake in home cages**

Regular chow intake over 4 h was measured daily from 11:00 am to 3:00 pm in their home cages. When the daily regular chow intake over 4 h was relatively stable for at least 3 successive days, mice were ready for food intake tests with drug treatment. On test days, regular chow or HFHS food intake was measured 45 min after IP drug infusion. In some experiments, mice were fasted for 24 h before food intake tests.

**Food intake test using feeding experimentation devices (FED) for meal pattern analysis**

FED, an open-source home-cage compatible device, was used for food intake test and operant behavior for food delivery(*5*). Mice were singly housed, and the FED device was placed in their cages for 6 days on a 12/12 on/off light cycle. For the free-feeding mode, a 20-mg pellet (Bio-Serv, NJ, USA) was dispensed into a feeding well of the FED device and monitored with a beam-break. When the pellet was removed, the time-stamp of removal, and the latency to retrieve the pellet were logged to the internal microSD card and a new pellet was dispensed. For fixed-ratio 1 (FR1) mode, FED logs the time-stamps of each "nose-poke" event to the internal storage. When the mouse activated the left nose-poke the FED delivered a combined auditory tone (4 kHz for 0.3s) and visual (all 8 LEDs light in blue) stimulus, and dispensed a pellet. While the pellet remained in the well both pokes remain inactive to prohibit multiple pellet delivery into the well. When the pellet was removed, the time-stamp of removal, and the latency to retrieve the pellet were logged. For chemogenetic manipulation of ZI DA neurons, saline or CNO (2.0 mg/kg) was injected at 19:00 immediately before dark cycle of the experimental day. Food intake was recorded for meal pattern analysis. For meal pattern analysis, a feeding bout was defined as a meal if ≥ 60 mg of food was ingested and if it was separated from another meal by ≥ 10 min(*33*).

**Dark/light conflict test with food**

Mice were exposed to HFHS food pellets 2 h daily, for at least 3 days, before food intake was tested in the light/dark box where HFHS pellets were placed in the light compartment. At the beginning of the test, mice were placed in the light compartment, facing the entry to the dark compartment. A digital camera over the light/dark box was used to track and record the activity of mice in the light compartment. Latency to first exit, total entries to the light compartment, percentage of time spent in the light compartment, food approaches, and food intake were recorded and measured in 10-min tests with or without photostimulation.

**Food intake in open-field chambers**

Mice were fasted for 24 hours prior to the test with free access to water. On the test day, all mice were placed into the same corner of an illuminated open field (50 cm x 50 cm x 38 cm) with an opaque, white acrylic floor. Food pellets were placed in a cup in the center of the open field and all mice were allowed to explore freely. Real-time activity of mice in the chambers were videotaped for data analysis using Ethovision XT video tracking and analysis software (Noldus, Wageningen, The Netherlands). Latencies to the first entry to the center area, total entries to the center area, and total time spent in the center area

(25 cm x 25 cm) were calculated with a limit of 10 minutes. After the tests, mice were removed from the open field and the numbers of pellet eaten were immediately counted.

**Operant lever presses for self-photostimulation**

At least one month after viral vector injection to induce ChIEF-tdTomato expression in ZI DA neurons of TH-Cre mice, mice were placed in an operant chamber (Med Associates, VT, USA) for self-photostimulation of PVT. Mice received photostimulation of 3 s immediately after every lever press. Continuous lever presses led to cumulative photostimulation. A digital camera was used to record the mouse movement track and behaviors during 30-min daily sessions for 9 days.

**Real-time place preference with photostimulation**

At least one month after viral vector injection to induce ChIEF-tdTomato expression in ZI DA neurons of TH-Cre mice, TH-Cre mice were tested for real-time place preference in a two-compartment chamber lacking additional contextual cues. One compartment was paired with a 20 Hz photostimulation and the other identical compartment was without photostimulation. Total test duration was 10 min.

**Conditioned place preference paired with photostimulation**

At least one month after viral vector injection to induce ChIEF-tdTomato expression in ZI DA neurons of TH-Cre mice, TH-Cre mice were first pretested for place preference in a two-compartment chamber with rough wire floor for one compartment and smooth floor for another compartment for 10 min. In the next 3 days following pretest, mice were trained for 10 min daily with photostimulation of PVT when they entered and stayed in the compartment with smooth floor. After photostimulation-paired training for 3 days, mice then received post-pairing place preference tests without photostimulation of 10 min daily for another 3 days. A digital camera was used to record the mouse movement track during the 10 min test daily.

**Conditioned place preference paired with HFHS food**

A 70 cm × 24 cm rectangular Plexiglass chamber with three compartments separated by removable Plexiglass walls was used for these experiments. The left and right compartments (28 × 24 cm each) had distinct wall patterns (black and white stripes versus white with black circles) and flooring (smooth versus rough wire floors). The center compartment measured 11.5 × 24 cm with no wall patterns and a smooth clear floor.

For acquisition of HFHS-associated place preference, mice first received pre-test for baseline preference. Subjects were placed in the center compartment and then the barriers were lifted that allowed the subject to explore the entire apparatus for 10 min. On days 2–5, two conditioning sessions were conducted per day, separated by at least 3 hours. In the first session mice were confined to one side compartment that contained a HFHS pellet for 30 min. Saline or CNO (2.0 mg/kg) was injected intraperitoneally 30-45 min before HFHS conditioning. In the second session, mice were confined to the opposite side compartment without food for another 30 min. On day 6, post-conditioning tests were conducted in the same manner as the baseline preference test.

For the expression of HFHS-associated place preference, mice also received pretest for baseline preference for 10 min similar to acquisition experiments above. On days 2-11, two conditioning sessions were conducted per day, separated by at least 3 hours. In the first session mice were confined to one side compartment that contained HFHS pellets for 30 min; in the second session, mice were confined to the opposite compartment without food for 30 min. HFHS-paired side was counterbalanced in each group. On day 12, post-conditioning tests were conducted in the same manner as baseline preference tests. 30 min before post-conditioning tests, saline or CNO (2.0 mg/kg) was injected intraperitoneally.

**In vivo fiber photometry**

Four weeks after virus injection and fiber optic implantation, a fiber photometry system (R810, RWD Life Science Co., Ltd, China) was used for recording fluorescence signal (GCaMP7f and isosbestic wavelengths) which was produced by an excitation laser beam from 470 nm and 410 nm LED lights. Calcium fluorescence signal were acquired at 60 Hz with alternating pluses of both lights. The power at the end of the optical fiber (200 μm, 0.37 NA) was adjusted to 30~40 μW. Recording parameters were set based on pilot studies that demonstrated the least amount of photobleaching, while allowing for the sufficient detection of the calcium response. A digital camera was used for behavioral recordings that were synchronized with calcium signal recordings.

To monitor the calcium signals of neurons associated with feeding behaviors, a FED3 device was used for the delivery of food pellet using a free-feeding mode. To minimize novelty-caused stress, mice were habituated for several days to the FED3 devices using the free-feeding mode. Before fiber photometry recordings, mice were fasted for 24 h to increase feeding motivation for the testing. On the experimental day, mice were allowed to acclimate in the home cage for 30 min with connection to the fiber photometry system. To better monitor the response of ZI DA and GABA neurons to food approach and consumption during a 30-min session, the FED3 device was set to deliver one 20-mg pellet (Bio-Serv, NJ, USA) 60 s after every previous pellet was retrieved by mice.

Regarding quantification, the filtered 410 nm signal was aligned with the 470 nm signal by using the least-square linear fit. ΔF/F was calculated according to (470 nm signal-fitted 410 nm signal)/ (fitted 410 nm signal). Normalized ΔF/F was obtained when the largest value of each trial was set to 100% and smallest value was defined as 0% using GraphPad Prism 9.

**Immunocytochemistry**

Mice were anesthetized with ketamine, and then perfused transcardially with saline followed by 4% paraformaldehyde or 3% glutaraldehyde. The 30-μm-thick coronal sections were cut on a cryostat, immersed in phosphate-buffered saline (PBS) for 1 h, treated with 2% normal horse serum in PBS for 1 h, and then incubated overnight at 4°C in polyclonal rabbit TH antibody (1:2000; AB152, Millipore) (*32, 34*) or anti-dopamine antiserum (1: 1000, from Dr. H. Steinbusch at Maastricht University) described in detail elsewhere(*35*). After washing in PBS, sections were placed in secondary Alexa488-conjugated (JacksonImmuoResearh, 711-545-152) or Alexa594-conjugated donkey anti-rabbit IgG (JacksonImmuoResearh, 711-585-152) at a dilution of 1:500 for 4 h, washed, and mounted on glass slides for imaging under microscope.

**Optogenetic stimulation**

Light was delivered through a fiber optic cable and into the brain of mice with a fiber optic implant using a blue-light laser (473 nm; Laserglow Technologies, Canada). Mice were briefly anesthetized using isoflurane to allow connection of the cable to the implant. Stimulation pulses (10 ms) of 20 Hz were controlled by a pulse generator (Doric Lenses, Canada) and delivered to stimulate ZI DA terminals in the PVT. During slice recordings, 20 Hz photostimulation was also delivered to target the PVT of brain tissues in the recording chamber.

##### Overall experimental design and statistical analysis

Animals were randomly assigned to different experimental groups before testing. Data collection and analysis were not performed blind to the conditions of the experiments. No statistical methods were used to predetermine sample sizes; our sample sizes are similar to those generally employed in the field. Data are expressed as mean ± s.e.m. Sample sizes (n number) refer to values obtained from individual animal in all behavioral experiments and individual cell recordings. Statistical analyses were performed using GraphPad Prism 9.0. Statistical significance was taken as *$p < 0.05$, **$p < 0.01$, ***$p < 0.001$ and ****$P < 0.0001$, as determined by one-way or two-way analysis of variance (ANOVA) followed by post hoc Bonferroni test, two-tailed paired or unpaired *t*-test.

#### Supplementary figures

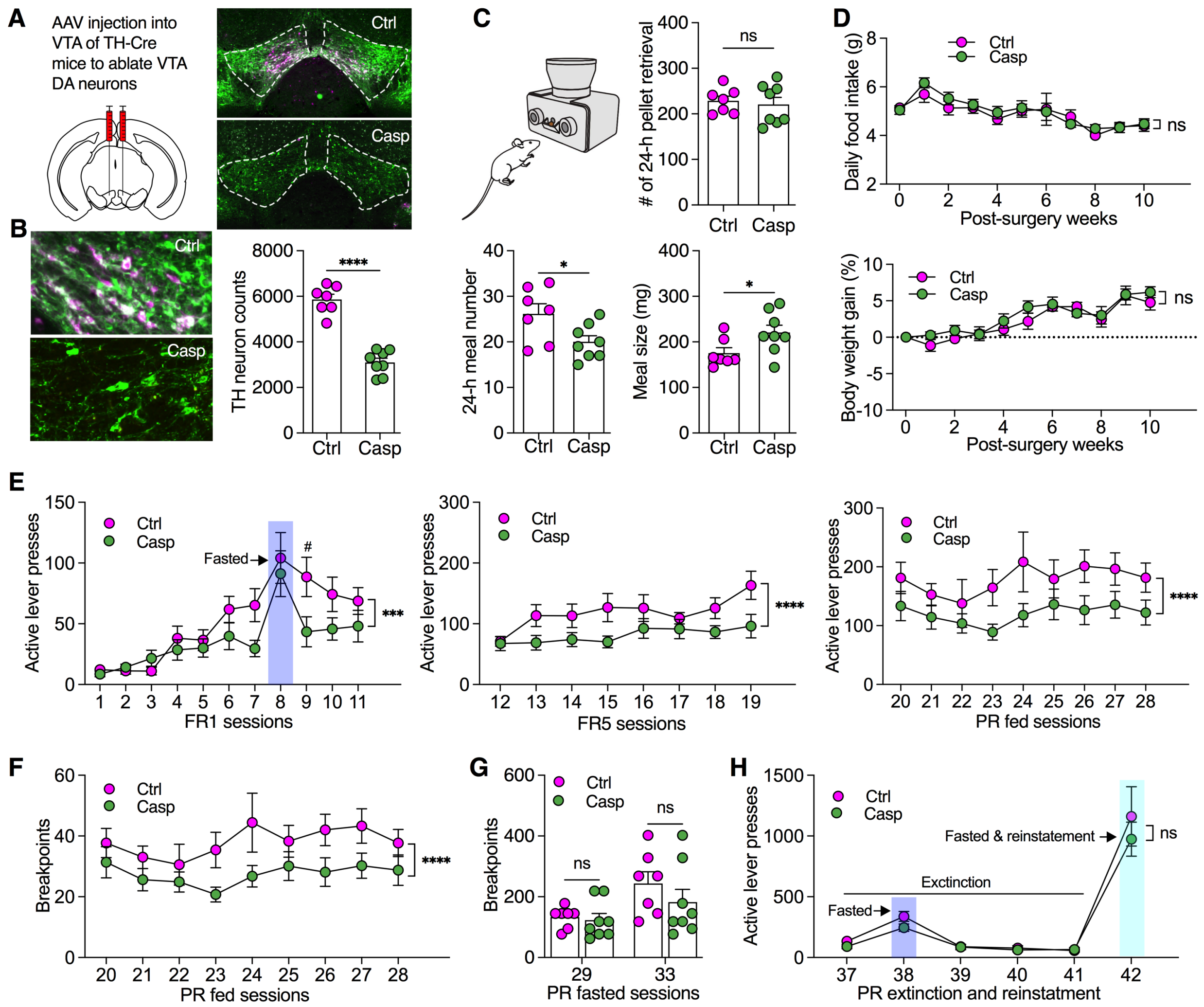


**Fig. S1: Selective ablation of VTA DA neurons reduced the motivational effort for seeking food reward at fed but not fasted states.** **A**, AAV was injected into VTA of TH-Cre mice to induce mCherry or mCherry plus caspase expression selectively in VTA DA neurons. **B**, TH-immunoreactive VTA neurons were ablated by virus-induced caspase expression. n= 7 mice for each group. Unpaired t test. **C**, Meal pattern analysis obtained from home-cage free feeding using FED devices shows total number of pellet retrieval, meal numbers, and averaged meal size of 24 h. n= 7 for ctrl and n= 8 for Casp group. Unpaired t test. **D**, Daily food intake and body weight gain for both groups (n= 7 for ctrl and n= 8 for Casp group) for 11 weeks following virus injection. **E**, Active lever presses of mice during FR1, FR5 and PR fed sessions of 45 min. Two-way ANOVA with Post hoc Bonferroni test. **F**, Breakpoint reached by fed mice (n= 7 for ctrl and n= 8 for Casp group) during operant PR sessions of 45 min. Two-way ANOVA with Post hoc Bonferroni test. **G**, Breakpoints reached by fasted mice (n= 7 for ctrl and n= 8 for Casp group). Two-way ANOVA with Post hoc Bonferroni test. **H**, Active lever presses at fed and fasted conditions (n= 7 for ctrl and n= 8 for Casp group) during PR extinction and reinstatement sessions. Two-way ANOVA with Post hoc Bonferroni test.

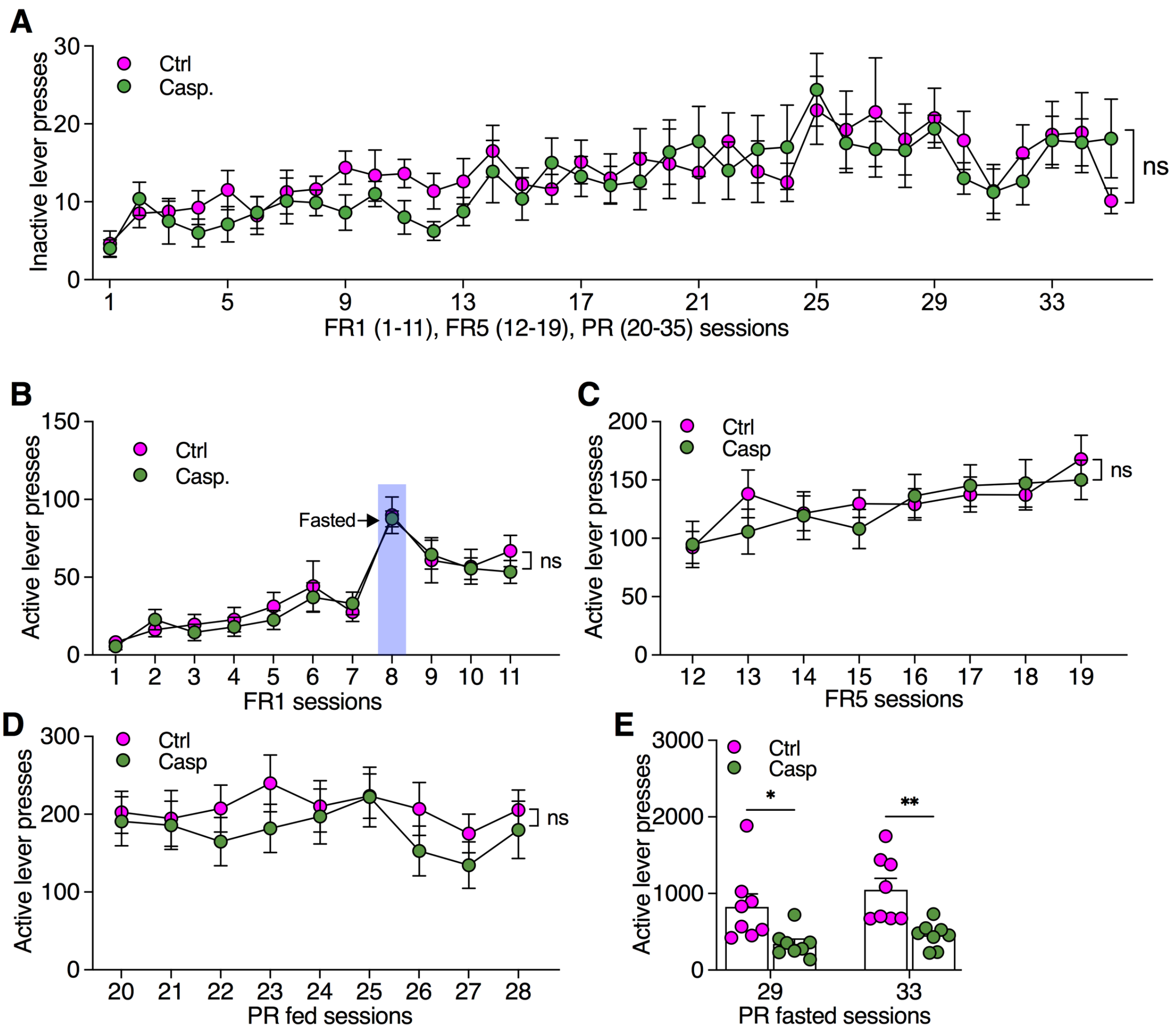


**Fig. S2: A**, Inactive lever presses of mice during FR1, FR5, and PR fed sessions of 45 min. Two-way ANOVA with Post hoc Bonferroni test. **B-E**, Active lever presses of mice during FR1, FR5, and PR fed and fasted sessions of 45 min. Two-way ANOVA with Post hoc Bonferroni test.

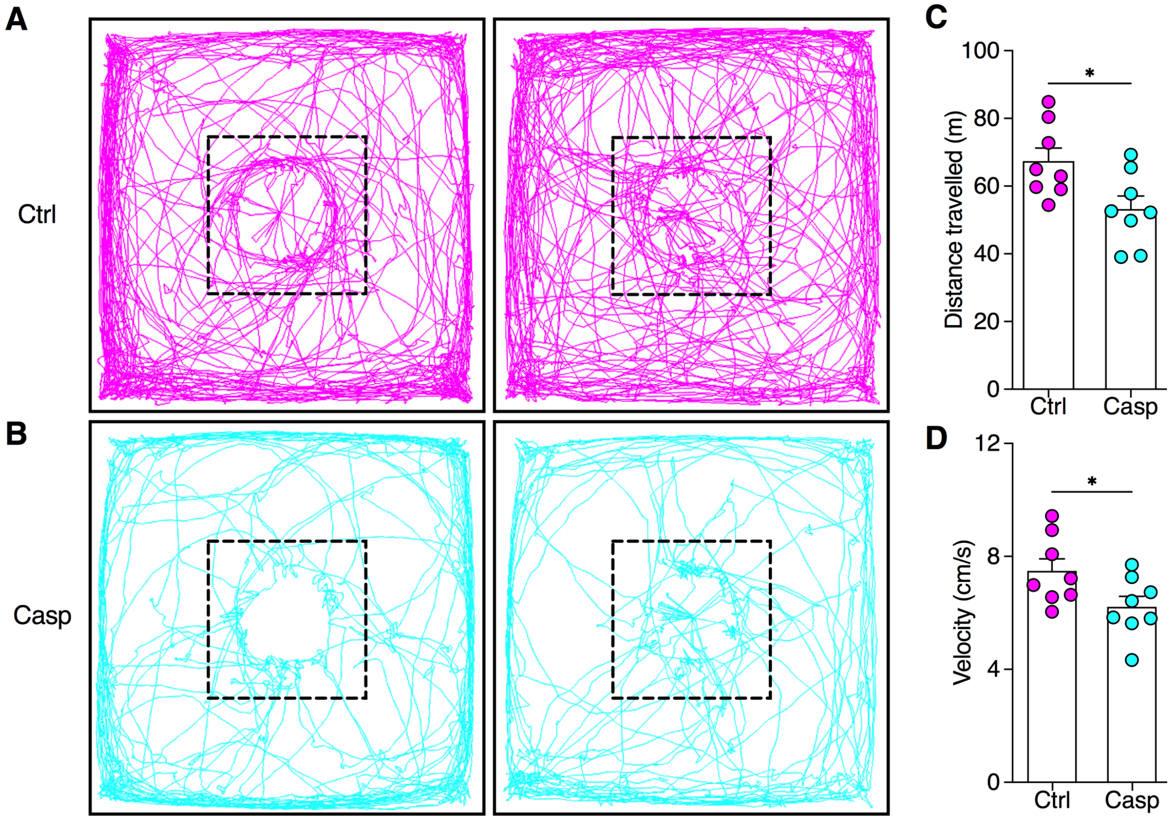


**Fig. S3: A,** Real-time activity of two control mice in open-field chambers with food pellets placed in the center. n= 7 mice each group. **B,** Real-time activity of two Casp mice in open-field chambers with food pellets placed in the center. **C**, Total of distance travelled in open-field chambers. n= 8 mice each group. Unpaired t test. **D**, velocity of mice travelled in open-field chambers. n= 8 mice each group. Unpaired t test.

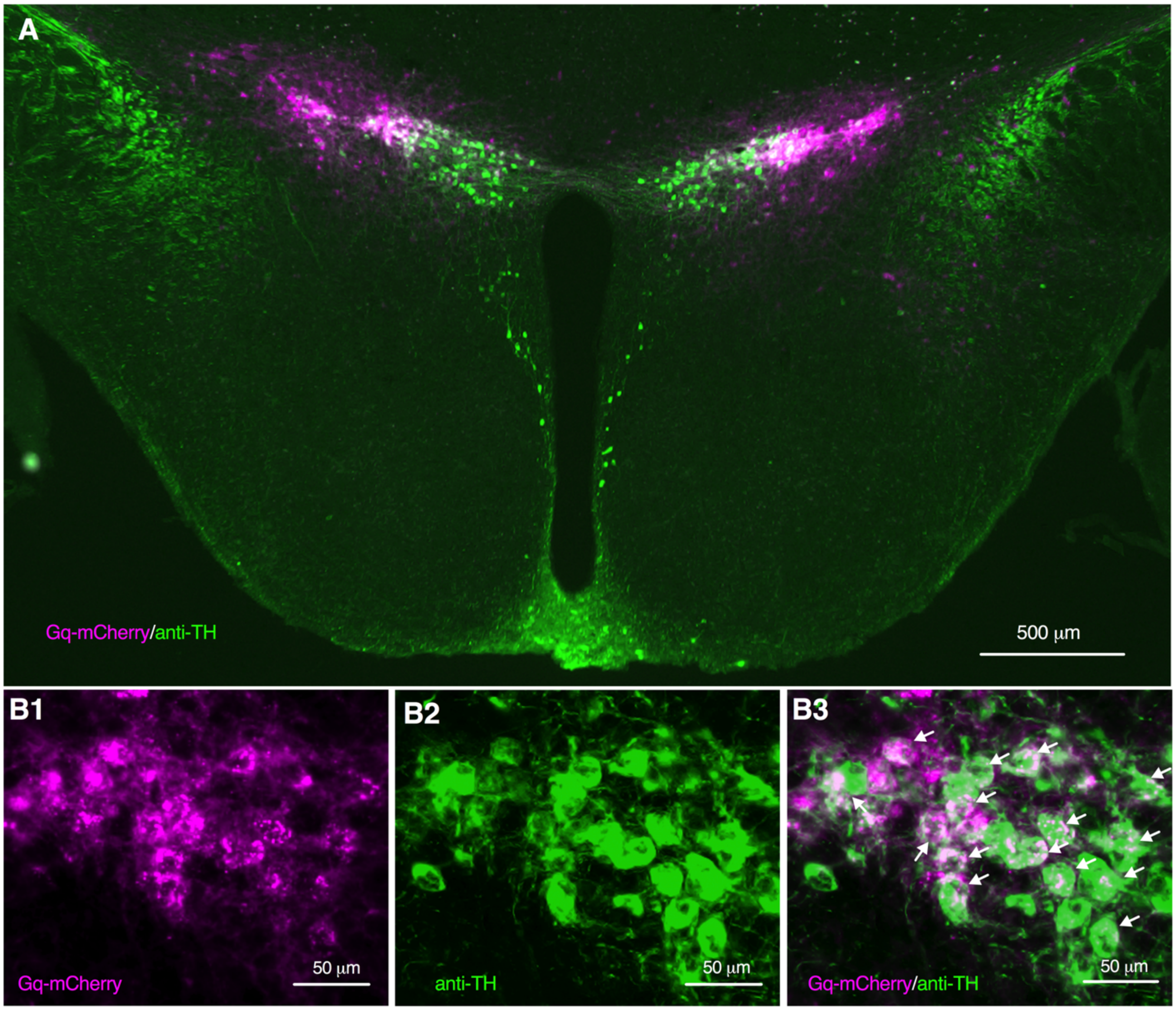


**Fig. S4: Excitatory DREADDS hM3D(Gq) was expressed in ZI TH neurons. A**, A fluorescent image showing both hM3D(Gq)-mCherry (pink) and anti-TH immunoreactivity (green) in ZI neurons of mice with viral injection. **B1-B3**, Zoomed-in images showing co-expression of hM3D(Gq)-mCherry and TH in ZI neurons.

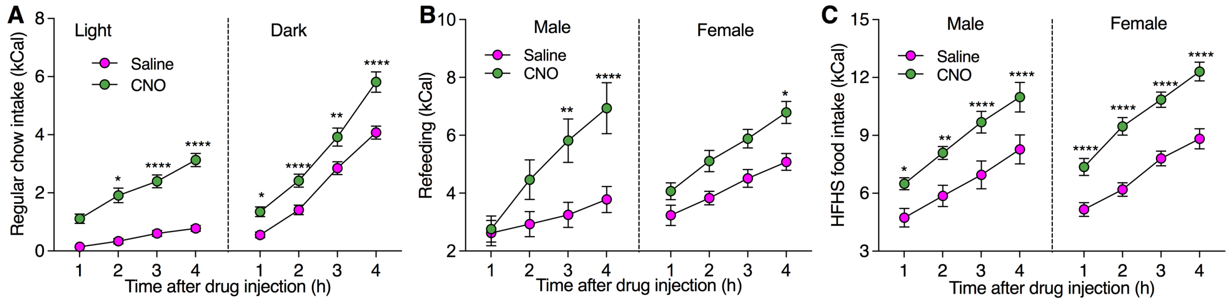


**Fig. S5: Chemogenetic activation of ZI DA neurons increased food intake of both male and female mice in different conditions. A,** Regular food in both light and dark cycles. **B**, Refeeding after 24 h fasting. **C**, HFHS food intake. Three-way RM ANOVA with Post hoc Bonferroni test.

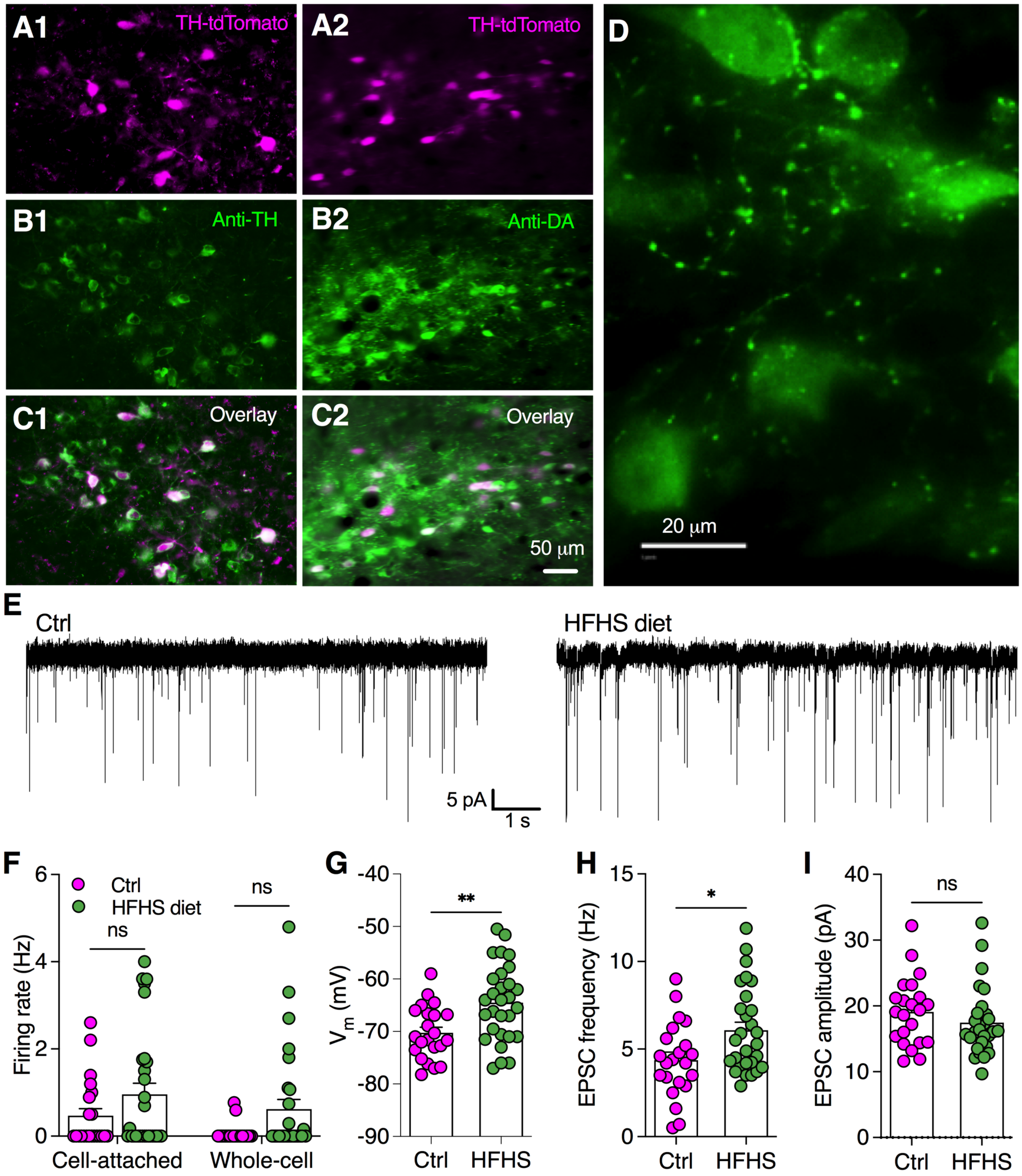


**Fig. S6: HFHS diet for 2 weeks potentiated excitatory synaptic transmissions onto ZI DA for the excitability of ZI DA neurons to increase the excitability of these neurons. A1-A3**, Representative images showing TH-immunoreactive neurons in ZI of TH-tdTomato mice. **B1-B3**, Representative images showing DA-immunoreactive neurons in ZI of TH-tdTomato mice. **C**, A zoomed-in image shows both DA-positive soma and axons in ZI. **D,** Representative traces showing the spontaneous EPSCs in ZI DA neurons from a control mouse and another mouse with HFHS diet. **E**, A bar graph with scattered data plots showing firing rate of ZI DA neurons from both control and HFHS-diet mice recorded in both cell-attached and whole-cell modes. Two-way ANOVA. **F**, Membrane potentials of ZI DA neurons from both control and HFHS-diet mice. Unpaired t test. **G**, sEPSC frequencies of ZI DA neurons from both control and HFHS-diet mice. Unpaired t test. **H**, sEPSC amplitudes of ZI DA neurons from both control and HFHS-diet mice. Unpaired t test.

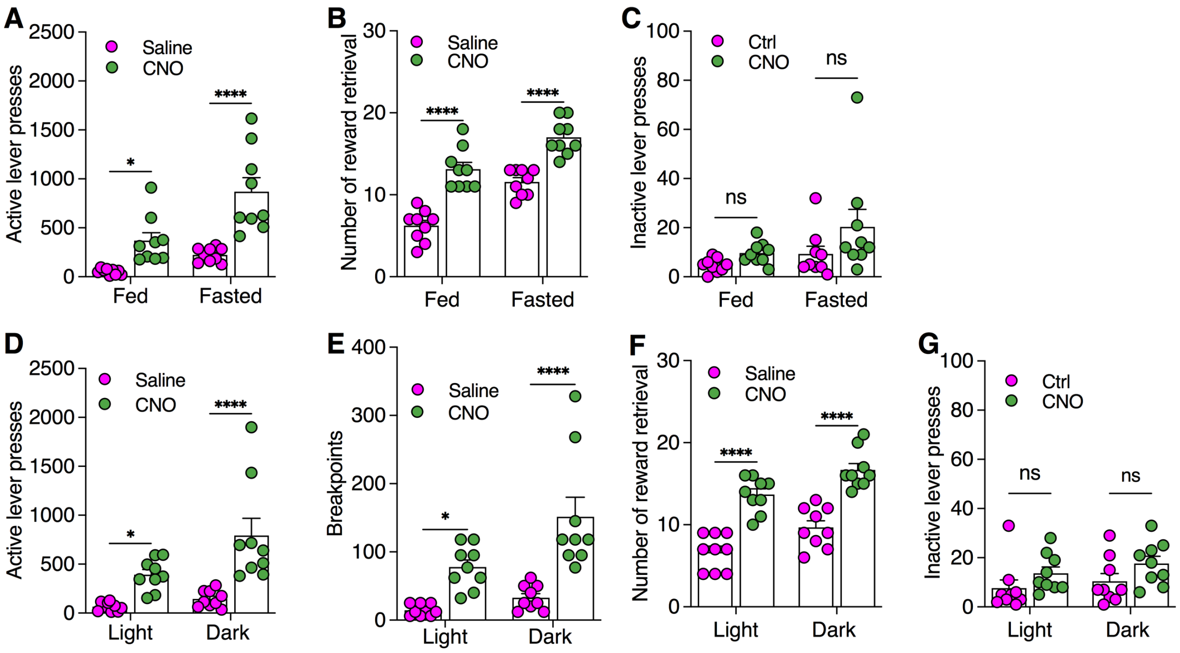


**Fig. S7: Chemogenetic activation of ZI DA neurons increased active lever presses for food reward but not inactive lever presses in both fed and fasted mice (A-C) during 45-min PR sessions with a stronger effect on tested in dark cycles (D-G).**

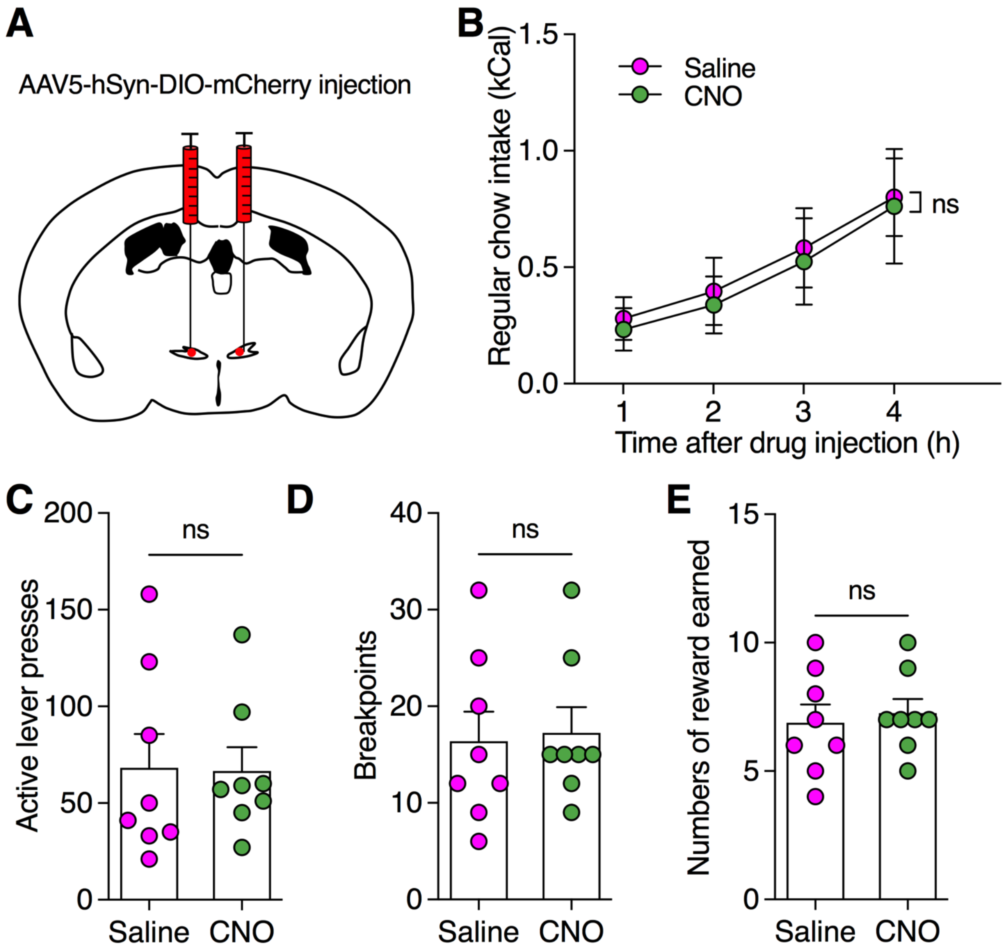


**Fig. S8: CNO injection had no effect on food intake or motivation for food rewards in mice with mCherry expression in ZI DA neurons. A**, A diagram showing AAV5-hSyn-DIO-mCherry was injected into bilateral ZI of TH-Cre mice. Two-way RM ANOVA with Post hoc Bonferroni test. **B**, IP injection of CNO had no effect on regular food intake over 4-h test following saline or CNO injection. **C-E**, IP injection of CNO produced no effect on active lever presses, breakpoints, or number of rewards earned during PR sessions of 45 min. Unpaired t test.

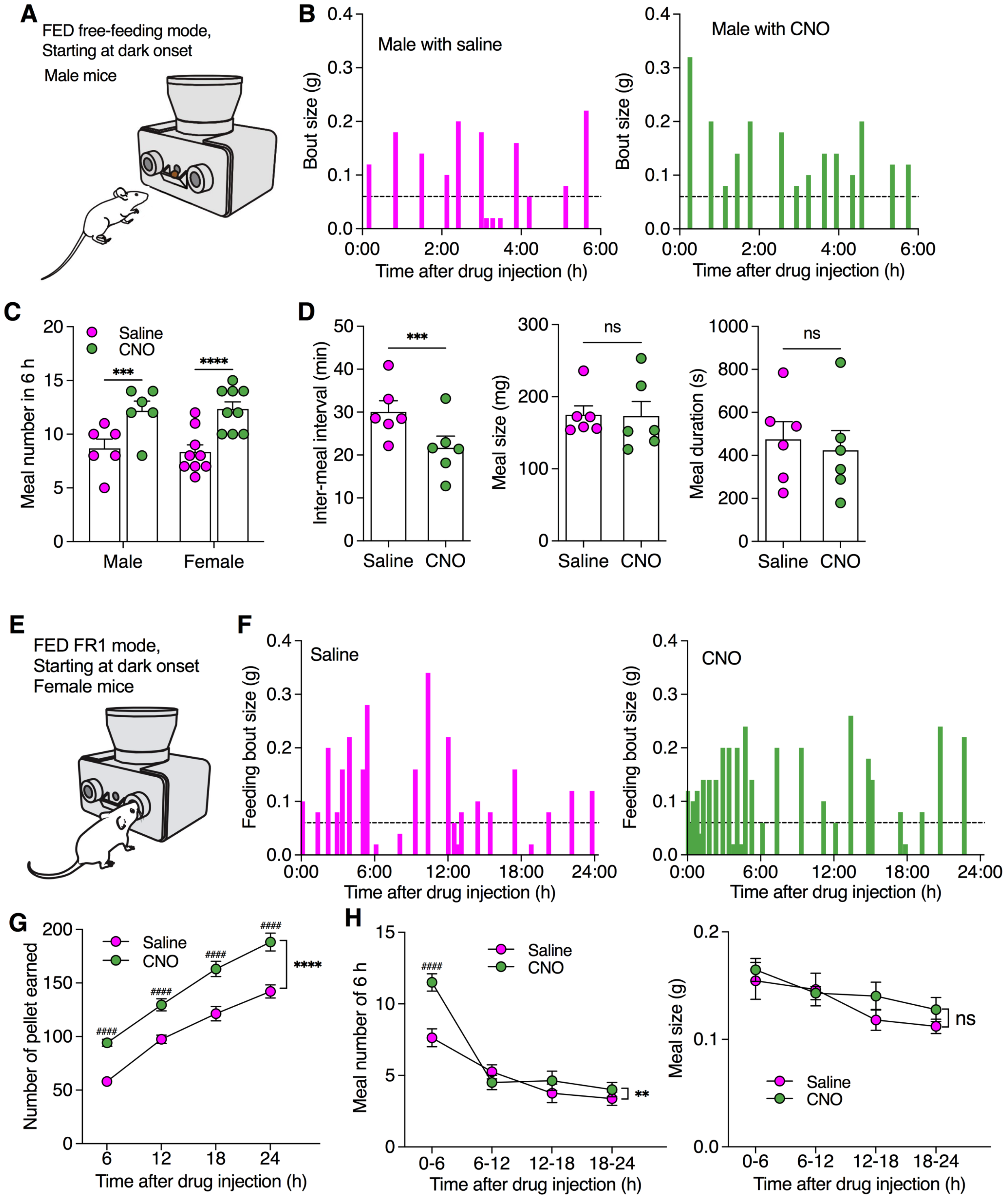


**Fig. S9: Chemogenetic activation of ZI DA neurons increased meal frequency but not meal size similarly in male and female mice tested with free-feeding and FR1 operant modes using FED devices. A,** A diagram showing FED device for free feeding test with regular food (20 mg per pellet) starting at dark onset. **B**, Bar graphs showing real-time feeding bout sizes following IP injection of saline and CNO (2.0 mg/kg) in a male TH-Cre mouse with hM3D(Gq)-mCherry expression in ZI DA neurons. **C**, Meal numbers of 6 h in both male (n= 6) and female (n= 9) mice following saline or CNO injection. Two-way RM ANOVA with Post hoc Bonferroni test. **D**, Inter-meal intervals (left), meal size (middle), and meal duration (right) following saline or CNO injection. n = 6 male mice each group. Paired t test. **E,** A diagram showing FED device under a FR1 mode with regular food (20 mg per pellet) starting at dark onset. One pellet was delivered immediately after each nose poke. **F**, Bar graphs showing real-time feeding bout sizes following IP injection of saline and CNO (2.0 mg/kg) in a male TH-Cre mouse with hM3D(Gq)-mCherry expression in ZI DA neurons. **G**, Number of pellets earned during a test duration of 24 h. n= 8 mice. Two-way RM ANOVA with Post hoc Bonferroni test. **H**, Meal number (left) and averaged meal size (right) per 6 h following IP injection of saline or CNO. n= 8 mice each group. Two-way RM ANOVA with Post hoc Bonferroni test.

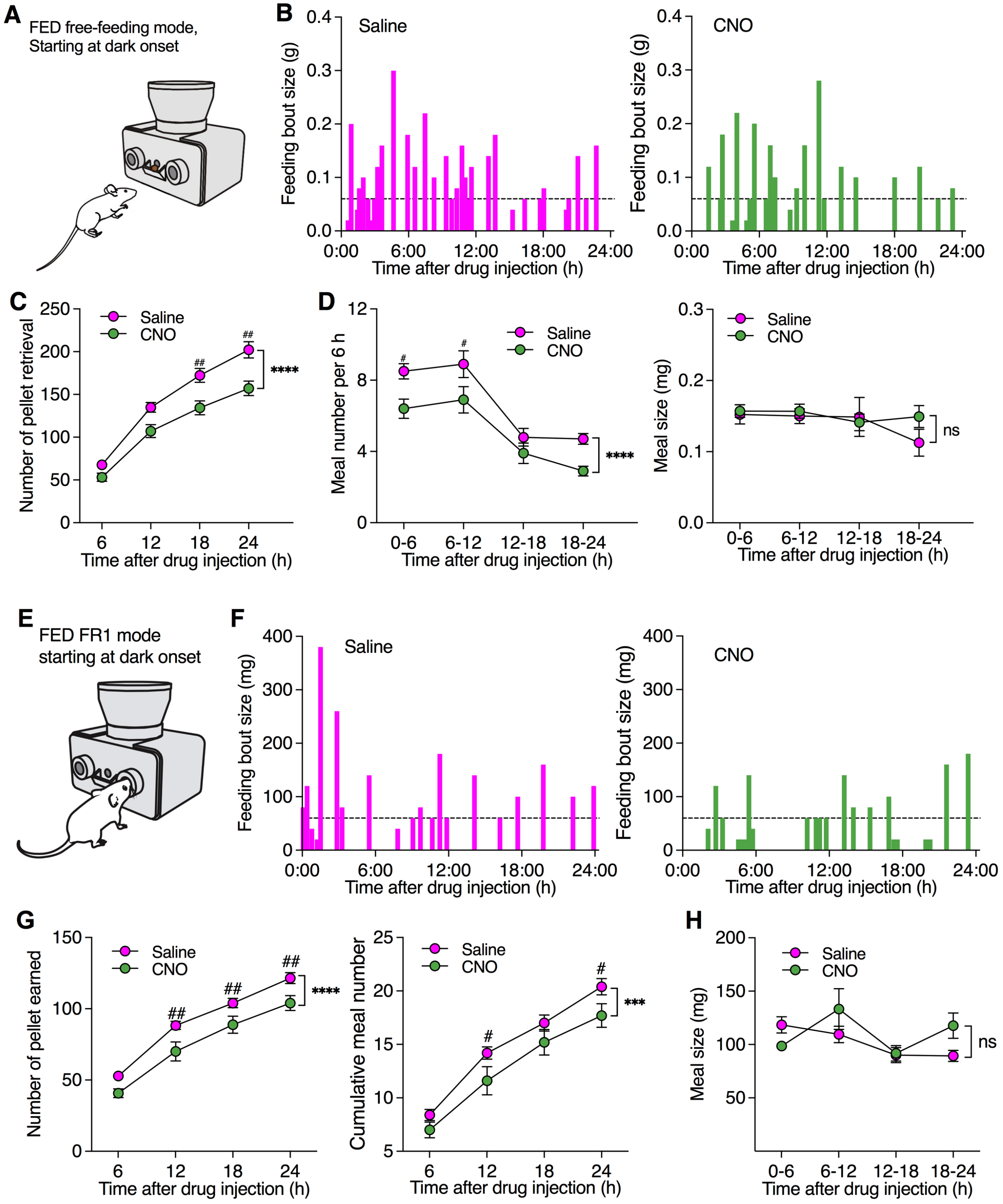


**Fig. S10: Chemogenetic inhibition of ZI DA neurons decreased meal frequency but not meal size tested with free-feeding and FR1 modes using FED devices. A,** A diagram showing FED device under free-feeding mode with regular food (20 mg per pellet) starting at dark onset. **B**, Bar graphs showing real-time feeding bout sizes following IP injection of saline and CNO (2.0 mg/kg) in a male TH-Cre mouse with hM4D(Gi)-mCherry expression in ZI DA neurons. **C**, Number of pellet retrieval during a test duration of 24 h. n= 10 mice. Two-way RM ANOVA with Post hoc Bonferroni test. **D**, Meal number (left) and averaged meal size (right) per 6 h following IP injection of saline or CNO. n= 10 mice each group. Two-way RM ANOVA with Post hoc Bonferroni test. **E,** A diagram showing FED device under a FR1 mode with regular food (20 mg per pellet) starting at dark onset. **F**, Bar graphs showing real-time feeding bout sizes following IP injection of saline and CNO (2.0 mg/kg) in a male TH-Cre mouse with hM4D(Gi)-mCherry expression in ZI DA neurons. **G**, Cumulative number of pellets earned (left) and meal number (right) over 24 h following IP injection of saline or CNO. n= 10 mice. Two-way RM ANOVA with Post hoc Bonferroni test. **H**, Averaged meal size (right) per 6 h following IP injection of saline or CNO. n= 10 mice each group. Two-way RM ANOVA with Post hoc Bonferroni test.

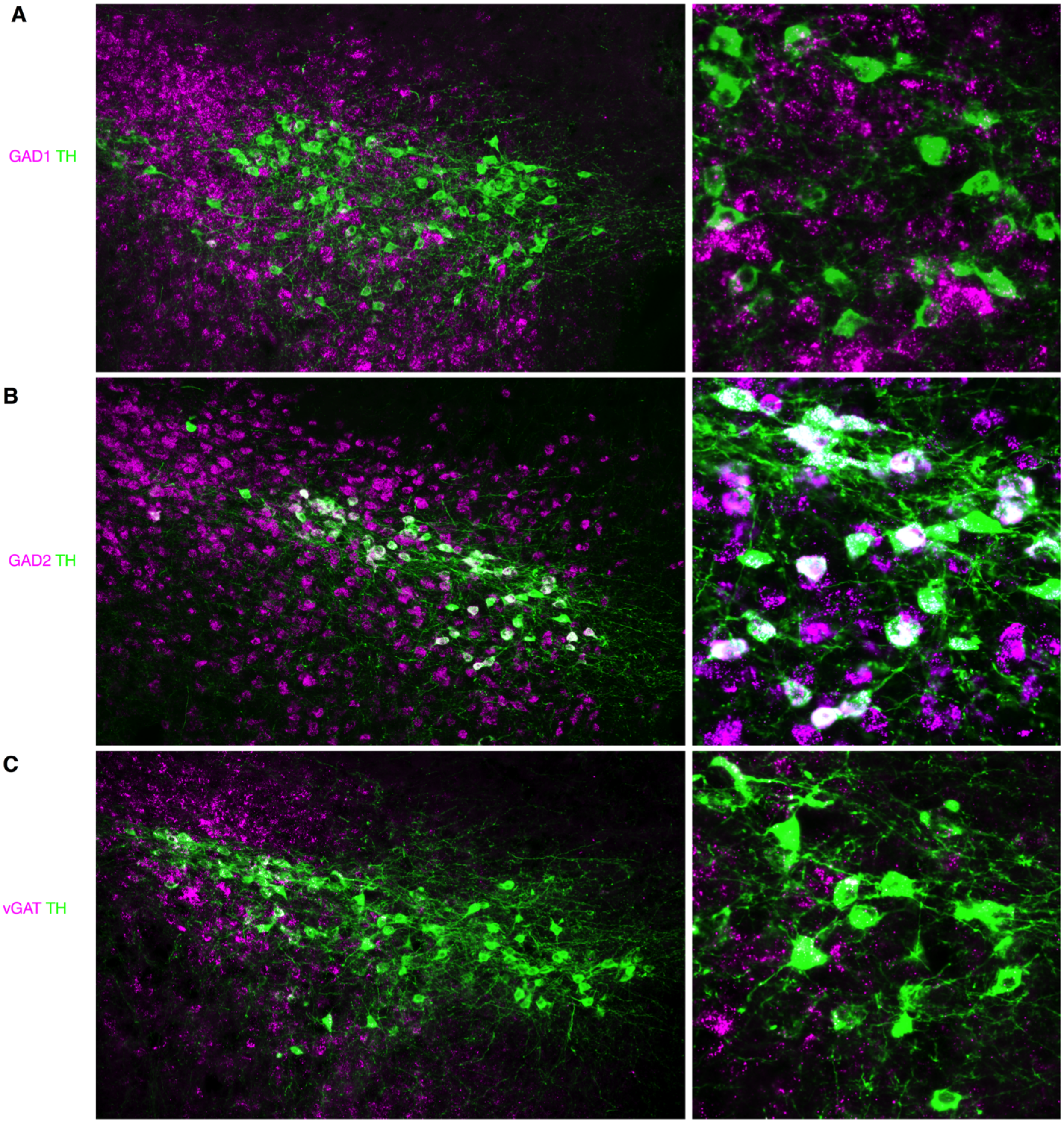


**Fig. S11: Majority of ZI DA neurons co-expressed GAD2 but not GAD1, and small population of them co-expressed vGAT.**

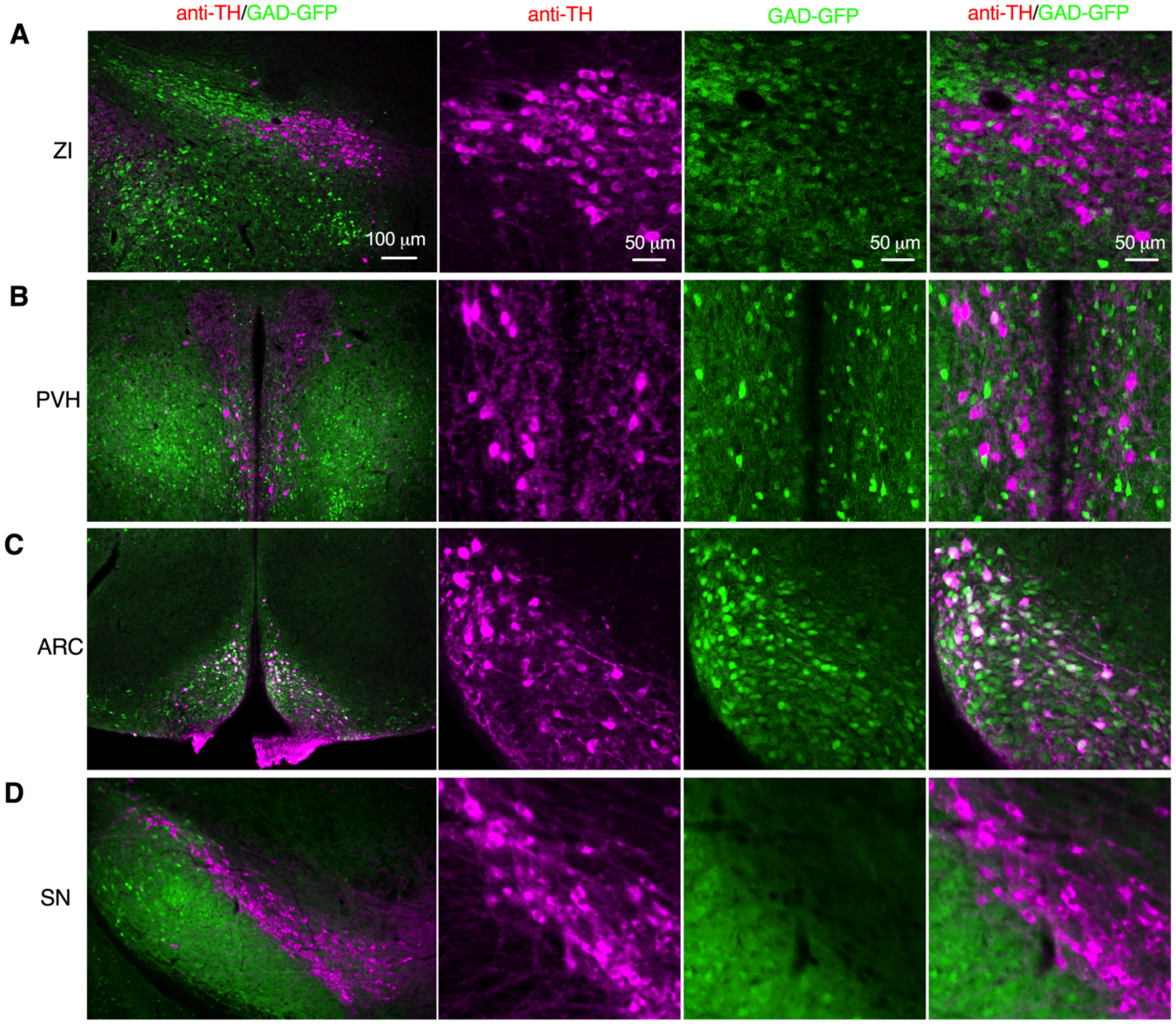


**Fig. S12: TH-immunoreactive neurons in ZI, paraventricular nucleus of hypothalamus (PVH), arcuate nucleus (ARC), and substantial nigra (SN) of transgenic GAG67-GFP mice.** Representative images showing no colocalization of GAD-GFP neurons and TH-immunoreactive cells in ZI (A) or SN (D), very low-level colocalization in PVH (B), but high-level colocalization in ARC (C).

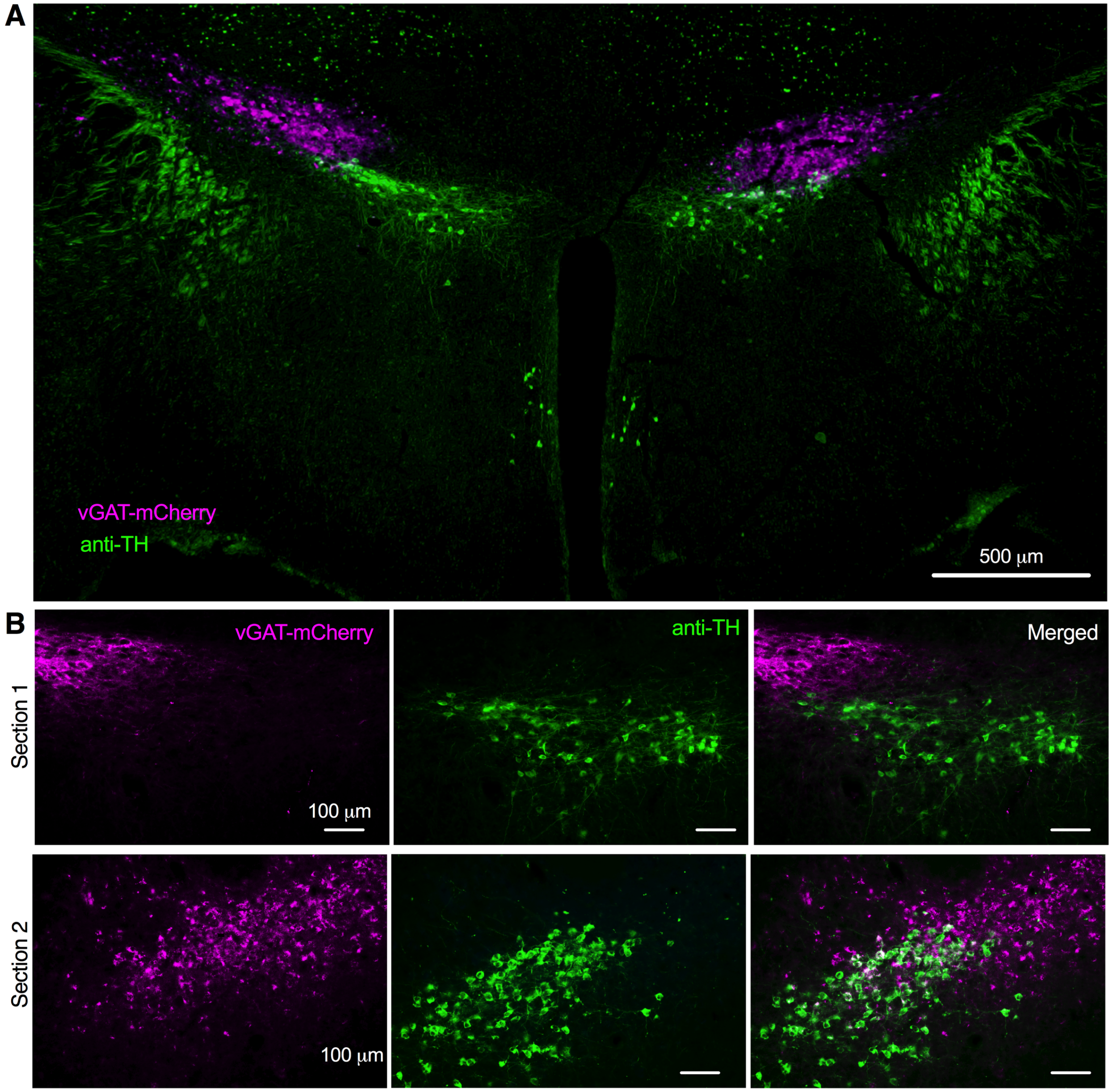


**Fig. S13: Very few hM3D(Gq)-mCherry-positive ZI GABA neurons were detected with TH-immunoreactivity in vGAT-Cre mice with ZI AAV-DIO-hM3D(Gq)-mCherry injection. A**, Representative image showing both ZI-vGAT-mCherry (purple) and anti-TH immunofluorescence (green) in ZI neurons. **B**, One representative coronal section showing no overlap of vGAT-mCherry and TH-immunoreactivity in ZI. **C,** Another coronal section showing high-density of both vGAT-mCherry and anti-TH immunoreactivity but little colocalization in ZI neurons.

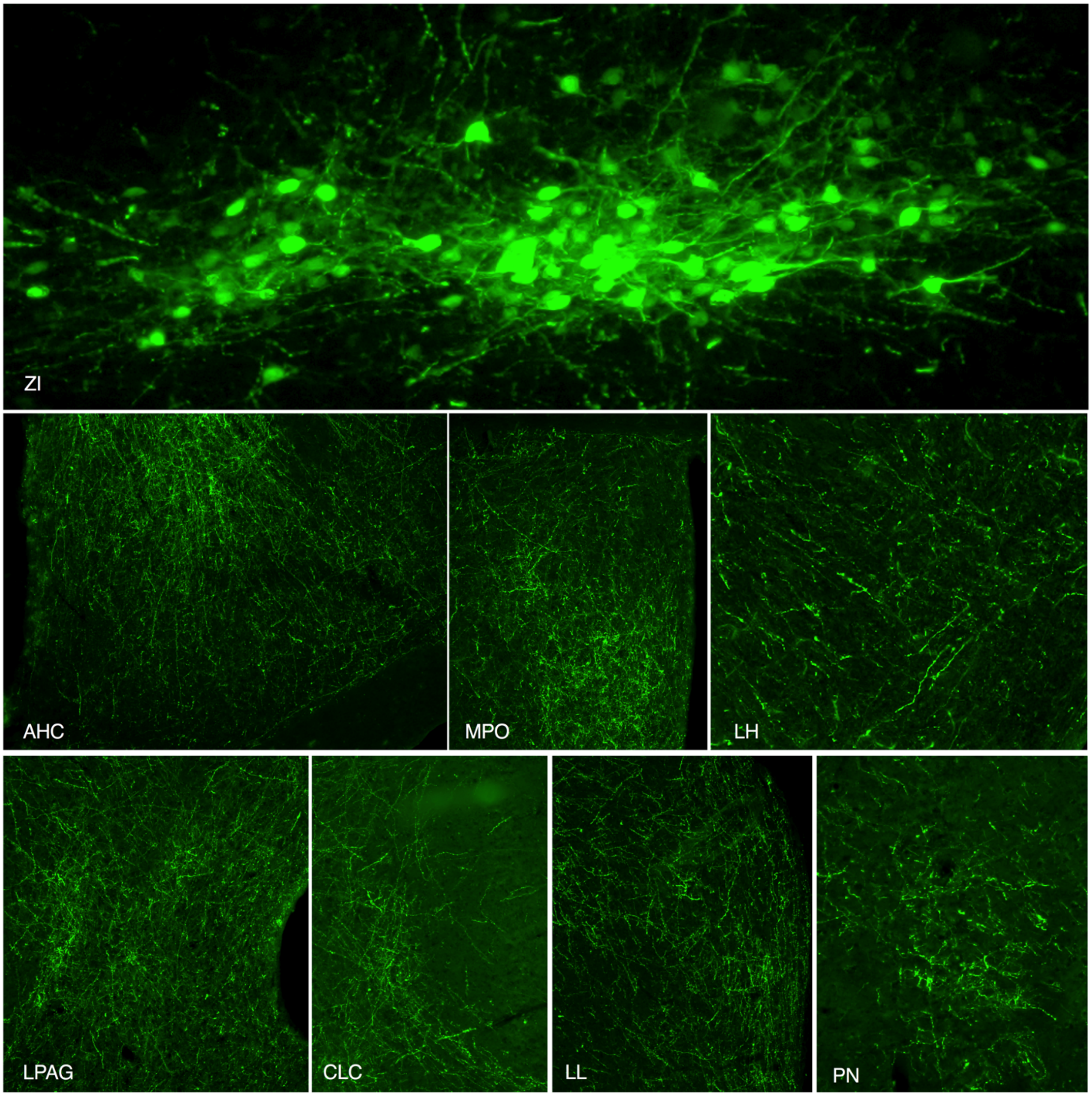


**Fig. S14: Other major brain regions targeted by ZI DA neurons.** AHC: anterior hypothalamus, central part**;** CLC: central nucleus of inferior colliculus; LH: lateral hypothalamus; LL: lateral lemniscus; LPAG: lateral periaqueductal gray; MPO: medial preoptic nucleus; PN: pontine nucleus; ZI: zona incerta.

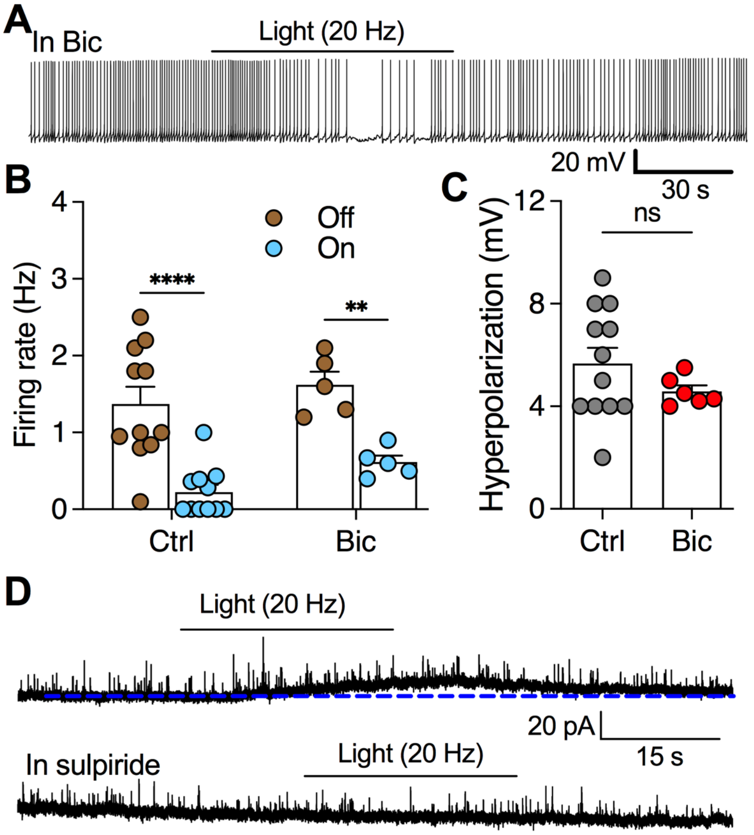


**Fig. S15: Photostimulation of ZI DA terminals in the PVT inhibited PVT neurons mainly by D2 receptor and partially by GABA$_A$ receptor activation. A,** A representative trace showing light-induced inhibition of PVT neuron innervated by ChR2-positive ZI DA terminals in the presence of Bic (30 μM). **B**. The firing rates of PVT neurons before and during photostimulation in the control condition and in the presence of Bic. **C**, The membrane hyperpolarization of PVT neurons in the control condition and in the presence of Bic. **D**, Representative traces showing a selective D2 receptor antagonist sulpiride (10 μM) abolished photostimulation-evoked tonic outward currents in PVT neurons.

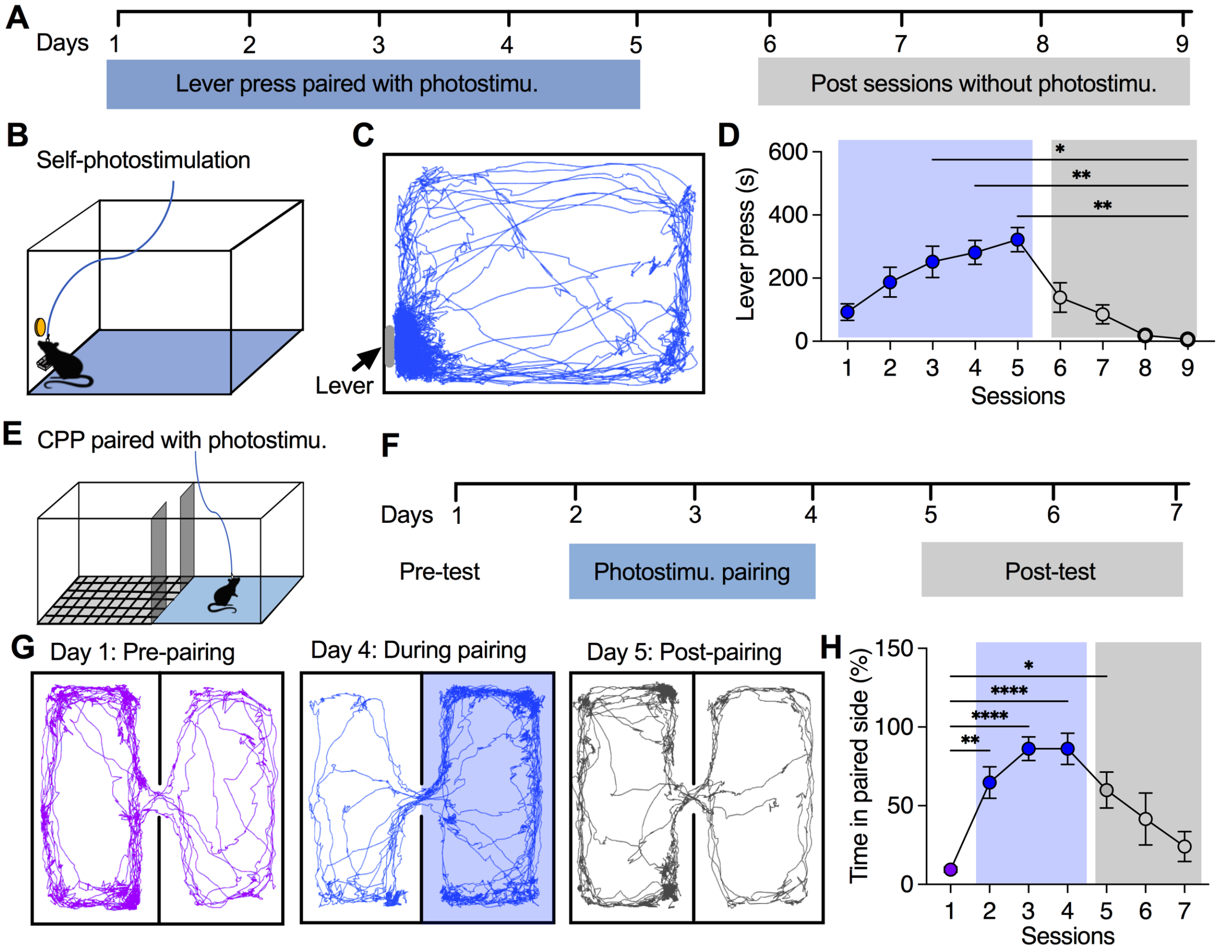


**Fig. S16: Activation of ZI-PVT DA projections evoked positive valence and promoted reward memory. A,** A schematic diagram showing the experimental process for self-photostimulation of ZI-PVT DA terminals. **B**, An operant chamber was used for self-photostimulation training and test. Mice with ChIEF expression in ZI DA neurons and fiber optics targeting PVT were used for the experiments. Photostimulation (20 Hz) was activated when lever was pressed down. **C**, A representative real-time motion track illustrates a mouse spent most of the time near the lever for evoking photostimulation. **D**, The cumulative lever-press time during a 30-min daily test session were recorded for 9 days. The first 5 sessions were paired with photostimulation of 20 Hz when the lever was pressed down, while the last 4 test sessions were paired with a sham stimulation. One-way RM ANOVA with Post hoc Bonferroni test. **E**, A diagram showing mice were placed in a two-compartment chamber for conditioned place preference test. One compartment was grounded with mess grids and another one remained smooth. The smooth compartmental side was paired with photostimulation (20 Hz). **F**, A schematic diagram showing the experimental process for pre-test without photostimulation, photostimulation-paired trainings, and post-test without photostimulation. **G**, Representative tracks show the real-time motion of a ZI-TH-tdTomato mouse in a two-compartment chamber before (day 1), during (day 4 of pairing), and post-paring test (day 5). **H**, Percentages of time that mice spent in compartmental side designed with photostimulation before, during and post photostimulation paring. n= 5 mice each group. One-way RM ANOVA with Post hoc Bonferroni test.

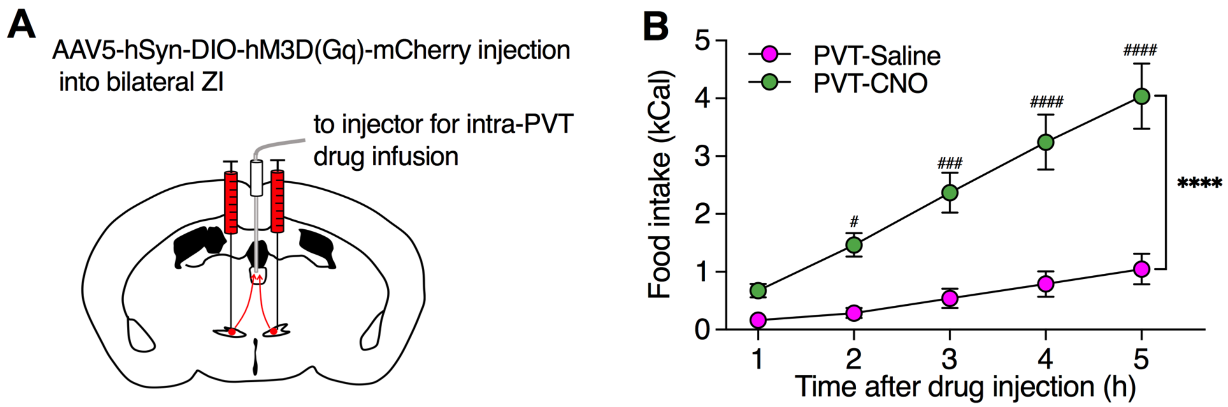


**Fig. S17: Chemogenetic activation of ZI DA axonal terminals in PVT increased food intake. A,** A diagram showing AAV5-hSyn-DIO-hM3D(Gq)-mCherry was injected into bilateral ZI of TH-Cre mice and a canular was implanted to target PVT. **B**, Food intake over 5 h after infusion of saline or CNO (0.5 mM, 0.5 μL) into PVT.

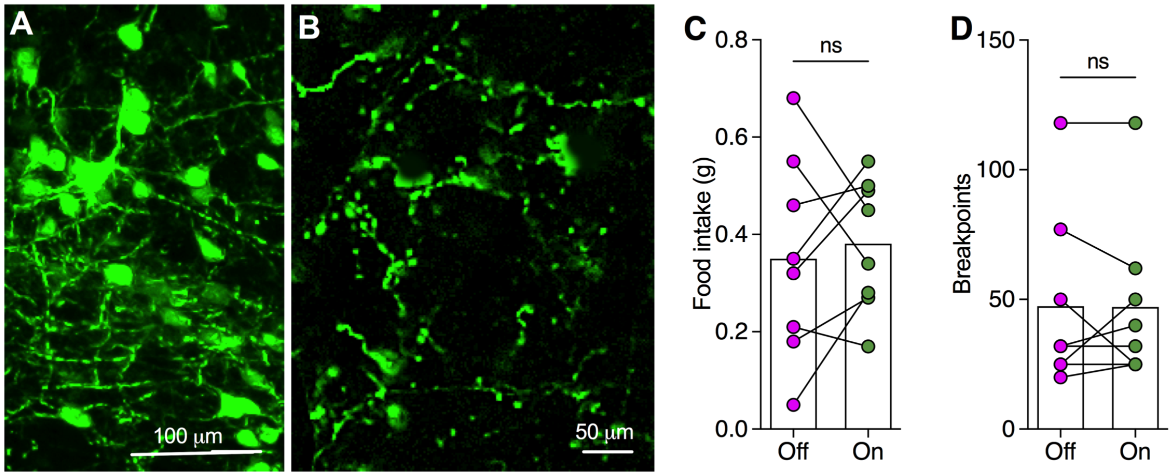


**Fig. S18: Optogenetic activation of ZI-PAG DA projections has no effect on food intake. A**, EYFP-positive DA neurons were found in ZI when AAV1-EF1a-DIO-ChR2(H134R)-EYFP-WPRE-HGHpA was injected into ZI of TH-Cre mice. **B**, EYFP-positive DA terminals were detected in PAG. **C**, Photostimulation (20 Hz) of ZI-PAG DA projections had no effect on HFHS intake. **D**, Photostimulation of ZI-PAG DA projections had no effect on breakpoints during PR sessions of 45 min.
